# Exacerbation of influenza virus induced lung injury by alveolar macrophages and its suppression by pyroptosis blockade in a human lung alveolus chip

**DOI:** 10.1101/2024.08.13.607799

**Authors:** Yuncheng Man, Yunhao Zhai, Amanda Jiang, Haiqing Bai, Aakanksha Gulati, Roberto Plebani, Robert J. Mannix, Gwenn E. Merry, Girija Goyal, Chaitra Belgur, Sean R. R. Hall, Donald E. Ingber

**Author notes:** Correspondence: Donald E. Ingber, M.D., Ph.D. Equal contribution (Yuncheng Man, Yunhao Zhai).

## Abstract

Alveolar macrophages (AMs) are the major sentinel immune cells in human alveoli and play a central role in eliciting host inflammatory responses upon distal lung viral infection. Here, we incorporated peripheral human monocyte-derived macrophages within a microfluidic human Lung Alveolus Chip that recreates the human alveolar-capillary interface under an air-liquid interface along with vascular flow to study how residential AMs contribute to the human pulmonary response to viral infection. When Lung Alveolus Chips that were cultured with macrophages were infected with influenza H3N2, there was a major reduction in viral titers compared to chips without macrophages; however, there was significantly greater inflammation and tissue injury. Pro-inflammatory cytokine levels, recruitment of immune cells circulating through the vascular channel, and expression of genes involved in myelocyte activation were all increased, and this was accompanied by reduced epithelial and endothelial cell viability and compromise of the alveolar tissue barrier. These effects were partially mediated through activation of pyroptosis in macrophages and release of pro-inflammatory mediators, such as interleukin (IL)-1β, and blocking pyroptosis *via* caspase-1 inhibition suppressed lung inflammation and injury on-chip. These findings demonstrate how integrating tissue resident immune cells within human Lung Alveolus Chip can identify potential new therapeutic targets and uncover cell and molecular mechanisms that contribute to the development of viral pneumonia and acute respiratory distress syndrome (ARDS).

## Introduction

Seasonal influenza A viruses (IAVs) cause around a billion cases of infection globally each year, including up to 5 million cases of severe illness and 650,000 cases of deaths^1^. IAVs remain a dangerous threat to the public heath as existing IAV strains may become highly transmissible and pathogenic reaching pandemic potential through rapid mutation and viral genome reassortment. Influenza H3N2 is one of the predominant IAV strains that infects alveolar epithelial cells, and it can cause severe viral pneumonia and acute respiratory distress syndrome (ARDS) characterized by high serum levels of pro-inflammatory cytokines and significant lung tissue injury^2, 3^. As current antiviral drugs are often susceptible to viral resistance and have limited ability to prevent virulent IAV-induced lung injury, it is imperative to identify novel therapies to combat future pandemics, and this can be accelerated using human-relevant preclinical models^4^.

Lung macrophages play major roles in maintaining lung homeostasis, host innate immunity, priming adaptive immunity, and tissue repairment and remodeling^5, 6^. A subset of lung residential macrophages strategically positioned within the alveolar space known as alveolar macrophages (AMs) serve as sentinel cells that recognize and phagocytose invading pathogens. They normally suppress local immune activation, but they also can drive a well-orchestrated, organ-level, inflammatory response if they fail to contain the pathogen^7, 8, 9^. These characteristics make them promising candidates for therapeutic treatment in lower respiratory tract infections. However, mounting evidence suggests that AMs originally derived from fetal progenitors are gradually outcompeted by bone marrow monocyte-differentiated macrophages in the lung as humans age^10, 11^, and these bone marrow-derived AMs exhibit greater immunoreactivity (e.g., enhanced glycolytic capacity and inflammatory gene expression) that can also lead to inflammation-associated tissue injury and severe outcomes in patients with respiratory viral infections^12, 13, 14^. But it is still poorly understood how these macrophages interact with lung parenchymal cells and microvascular endothelial cells (ECs) and cause lung injury after exposure to viruses.

It is difficult, if not impossible, to define the contribution of a specific cell type to complex physiological responses in vivo, either in animal or in human clinical studies, and key lung features that may be involved in the host injury response to viral infection (e.g., alveolar-capillary tissue barrier, physiologic mechanical cues associated with breathing motions) are missing from conventional *in vitro* static cultures as well as more sophisticated lung organoid models^15^. However, human organ-on-a-chip (Organ Chip) microfluidic culture models of the lung alveolus incorporate these features and permit experimentalists to vary cell type composition and other key control parameters (e.g., breathing motions, chemical factors, pathogens, etc.) independently^4, 16, 17, 18^. Thus, in the present study, we leveraged a human Lung Alveolus Chip that was previously used successfully to model influenza H3N2 virus infection, as well as drug toxicity-induced lung edema, intravascular thrombosis, lung cancer, and radiation-induced lung injury^4, 19, 20, 21, 22, 23^. However, we modified the model by incorporating resident human AMs to add an additional level of organ-level mimicry that has not been explored in the past in order to investigate how bone marrow-derived AMs contribute to IAV pathogenesis and lung injury. We also carried out these studies in the absence of breathing motions to focus specifically on how AMs interplay with pulmonary epithelium and endothelium during the response to viral infection because past work has shown that mechanical forces can act directly on macrophages to suppress inflammation^24^.

## Results

### Resident macrophages reduce influenza virus replication in the human Lung Alveolus Chip

The human Lung Alveolus Chip is a microfluidic device that contains two parallel channels separated by an ECM-coated porous membrane lined with primary human lung alveolar epithelial cells on the apical side and lung microvascular ECs on the basal side, mimicking the alveolar-capillary interface of human alveoli (**Fig. 1a**). The epithelial cells are cultured under an air-liquid interface while the endothelium is exposed to continuous medium flow to mimic vascular perfusion. To model bone marrow-derived AMs that reside within the alveolar space, CD14^+^ monocytes from peripheral human blood were differentiated into macrophages using human macrophage-colony stimulating factor (M-CSF) and interleukin (IL)-4, and then cultured on top of the alveolar epithelium within the apical channel. Flow cytometric analysis of two macrophage surface markers, CD163 and CD209^25^, confirmed that over 90% of cells were mature macrophages following differentiation (**Supplementary** Fig. 1a). Confocal imaging of the distribution of epithelial and endothelial cell-cell adhesion proteins (E-cadherin and VE-cadherin, respectively), along with CD45 stained macrophages, confirmed that the macrophages spontaneously positioned themselves as resident AMs adherent to the apical surface of the alveolar epithelium, which form a tight interface with the endothelium below under baseline conditions (**Fig. 1b**). Flow cytometry analysis of cells isolated from the apical channel using CD326 (EpCAM) staining for alveolar epithelial cells and CD45 for macrophages confirmed that there was a low ratio of macrophages to alveolar epithelial cells (∼1:12; **Supplementary** Fig. 1b), which is consistent with human lung alveolar physiology^26, 27^.

**Fig. 1.**
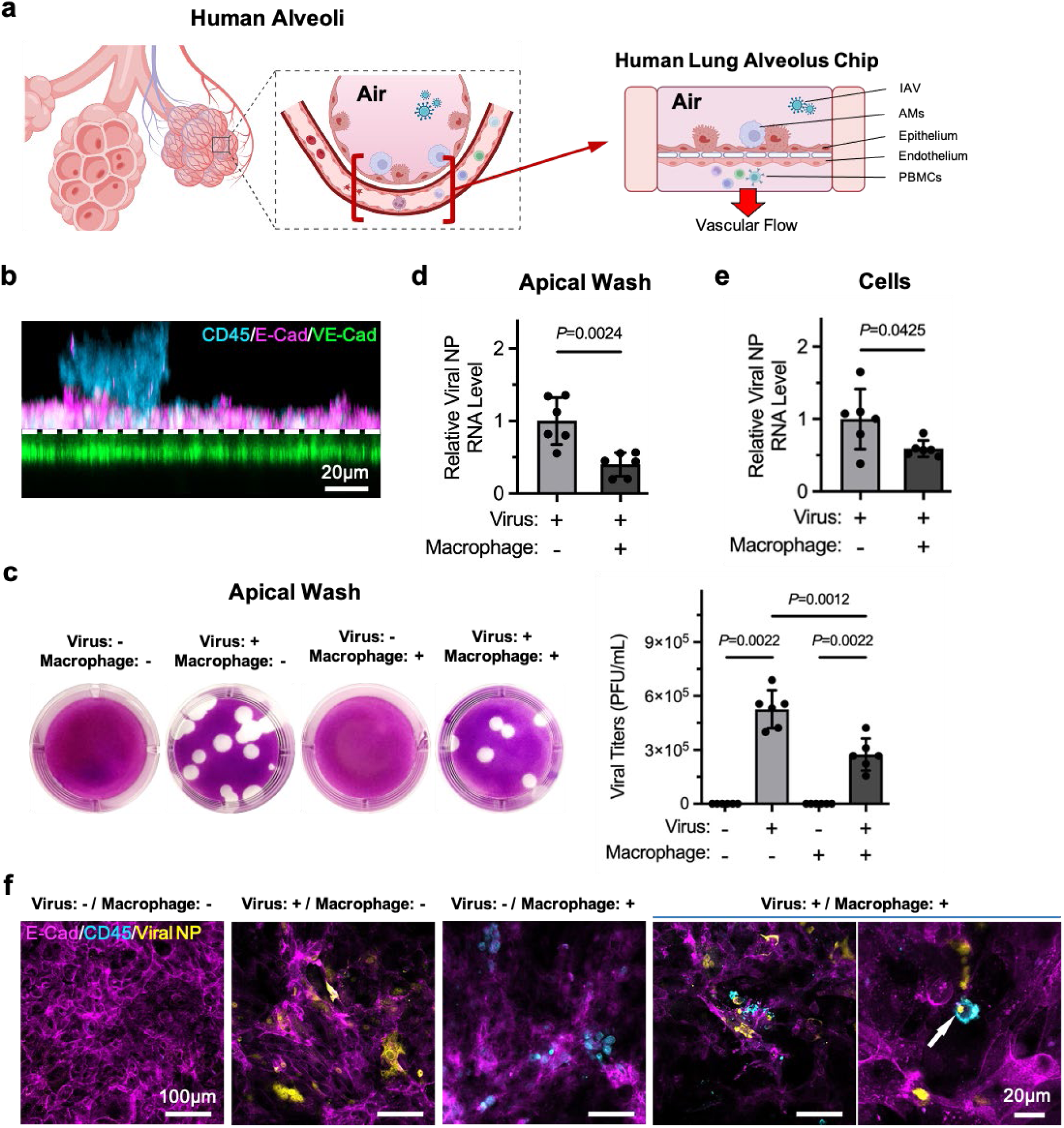
The human Lung Alveolus Chip model of influenza A virus infection and direct observation of the role of macrophages after infection. a Schematic of the human lung alveolus with IAV infection (created with BioRender.com), showing type I and II alveolar epithelial cells, alveolar macrophages, lung microvascular endothelial cells and circulating immune cells. Schematic of human Lung Alveolus Chip with IAV infection is also shown, featuring recapitulation of key cellular components, tissue barrier and physiologic vascular flow. IAV: influenza A virus. AMs: alveolar macrophages. PBMCs: peripheral blood mononuclear cell. b Z-stack confocal imaging showing the triple-culture human Lung Alveolus Chip model. c Images of plaques (left) for quantifying viral titers in the apical wash samples (right) collected from infected or mock infected chips, with or without macrophages. d RT-qPCR analysis of relative viral NP RNA levels in the apical wash samples collected from infected chips with or without macrophages. e RT-qPCR analysis of relative viral NP RNA levels in the total RNA extracted from the apical channels in infected chips with or without macrophages. f Immunofluorescence showing positive IAV infection of the alveolus chip with or without macrophages at 48 h post infection with A/Hong Kong/8/68 (H3N2) viruses. Macrophage activities after infection were shown by two magnifications of microscopic images. The white arrow indicates macrophage phagocytotic activity. Data shown are mean ± SD; n = 6 chips in each group in c, d and e; unpaired t-test.

The modular nature of the Lung Alveolus Chip enabled construction of chips with or without AMs in the alveolar space of the apical channel, allowing us to interrogate their contribution to the human lung response to viral infection. Lung Alveolus Chips were cultured with or without macrophages, and were then infected with a pandemic influenza H3N2 virus strain (A/Hong Kong/8/68) at a multiplicity of infection (MOI) of ∼4 (approximately 50,000 epithelial cells per chip) or mock infected (adding fresh culture medium without virus). The experimental timeline (**Supplementary** Fig. 1c) involves culturing the chips for 11 days, with the last 4 days incorporating an ALI above the epithelium on-chip to ensure optimal differentiation, before the addition of macrophages and subsequent exposure to IAV. Robust viral infectivity was detected on-chip in the absence of macrophages at 48 h post infection when apical effluents were analyzed using plaque assays, as previously reported^4^, and there was no detectable virus in effluents from control uninfected chips without macrophages (**Fig. 1c**). We also detected viral titers in the Lung Alveolus Chips with macrophages and not in uninfected chips; however, the viral load was significantly reduced by approximately 50% (**Fig. 1c**). Reverse transcription-quantitative polymerase chain reaction (RT-qPCR) analysis independently confirmed that both the RNA levels for influenza viral nucleoprotein (NP) in the apical effluents and in the total RNA extracted from the cells were again significantly reduced by about one half in infected chips that contained macrophages compared to those without (**Fig. 1d, e**). Importantly, these results replicate *in vivo* findings where AMs have been shown to protect against IAV infection by limiting viral spreading^28, 29^.

Immunofluorescence carried out in parallel confirmed successful viral infection on-chip as indicated by positive cellular expression of NP (**Fig. 1f**). While the alveolar epithelium was lined by cells surrounded by continuous E-cadherin containing cell-cell adhesions in non-infected chips, infection with influenza H3N2 resulted in disruption of these cell junctions and reduced E-cadherin staining by 48 h post infection (**Fig. 1f**). While most macrophages remained intact in control uninfected chips at 2 days after IAV infection, many of these CD45^+^ cells appeared non-refractile or broken into pieces, suggesting induction of significant macrophage cell death (**Fig. 1f**). Interestingly, in some field of views (FOVs), we observed colocalization of intact CD45^+^ cells and viral NP (**Fig. 1f**), suggestive of phagocytosis of infected epithelial cells by these macrophages, which is again consistent with the reduction in viral load observed in these chips. Altogether, these results indicate that the human Lung Alveolus Chip recapitulates many features of human alveolar infection by IAVs where AMs initially fight off the virus but sustained viral infection can induce massive loss of AMs, cell injury, and disruption of alveolar epithelial cell-cell junctions leading to compromise of pulmonary barrier function and ARDS^10, 30^.

### Macrophages exacerbate alveolar tissue injury after influenza infection

As monocyte-derived AMs in human lung have been shown to determine disease severity in patients with influenza,^11, 14^ we next sought to use the Lung Alveolus Chip to further interrogate their role in IAV pathogenesis. Analysis of pro-inflammatory cytokine secretion into the apical wash of the chips 48 h post infection revealed that viral infection induced significant upregulation of IL-6, IL-8, TNF-α, IP-10, and RANTES, but not IL-1β, IL-18, macrophage inflammatory protein (MIP)-1α, or monocyte chemoattractant protein (MCP)-1 when macrophages were not present (**Fig. 2a**). Importantly, when we co-cultured macrophages in the chips, we found that in addition to these same pro-inflammatory cytokines, IL-1β, IL-18, MIP-1α, and MCP-1 were also significantly upregulated (**Fig. 2a**). Moreover, the levels of these pro-inflammatory cytokines were increased in infected chips with macrophages compared to those without, consistent with macrophage-exacerbated lung inflammation.

**Fig. 2.**
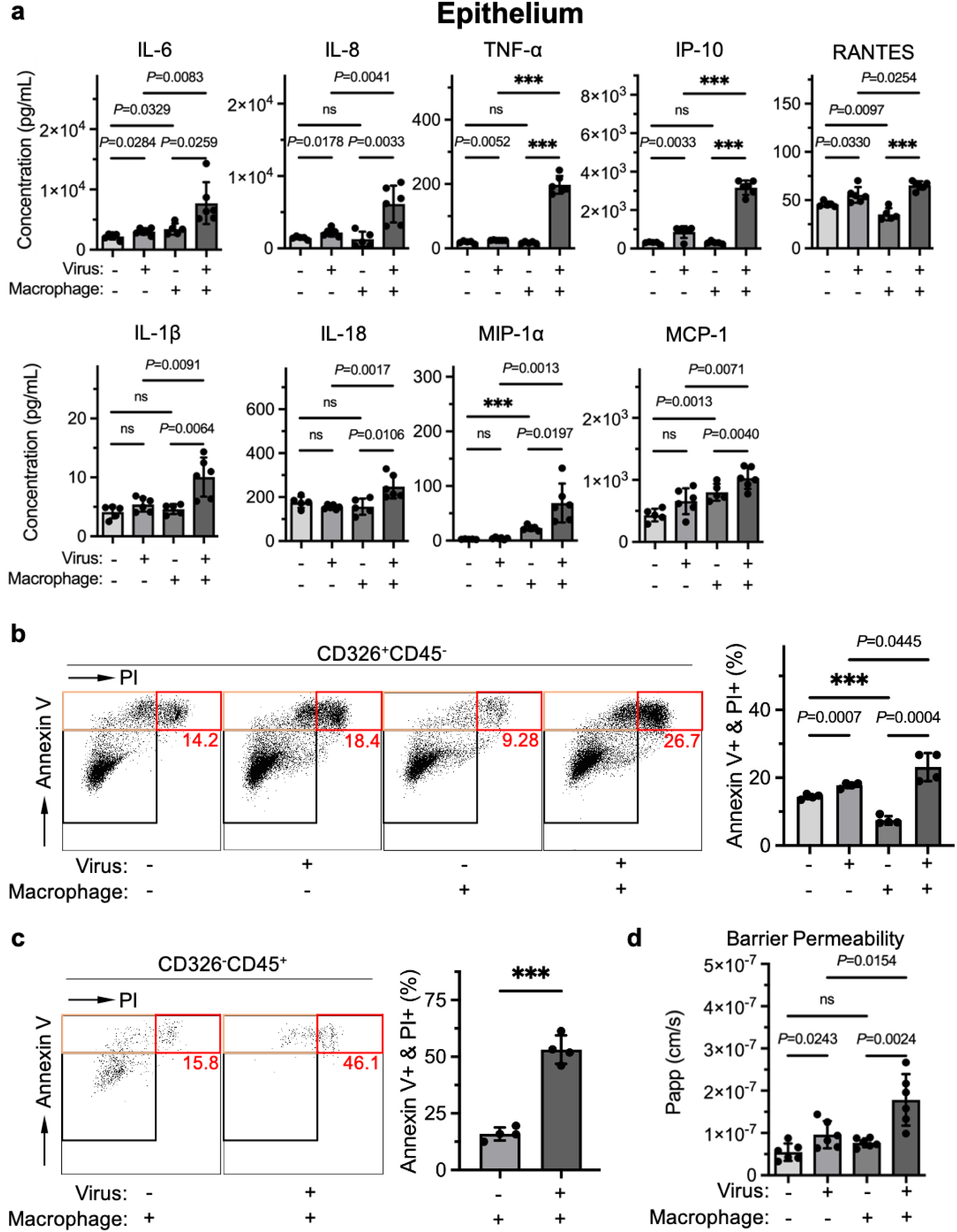
Macrophages exacerbate lung alveolar tissue inflammation and injury after influenza infection in the alveolus chip. **a** Production of the indicated cytokines at 48 h post infection measured by Luminex assay in the apical wash samples collected from infected or mock infected chips, with or without macrophages. **b** Dot plots by flow cytometry analysis (left) for quantifying alveolar epithelial cell deaths (Annexin V^+^/PI^+^) (right) in infected or mock infected chips, with or without macrophages, at 48 h post infection. **c** Dot plots by flow cytometry analysis (left) for quantifying macrophage deaths (Annexin V^+^/PI^+^) (right) in infected or mock infected chips at 48 h post infection. **d** Barrier function assay showing macrophage-exacerbated alveolar-capillary tissue barrier permeability after infection. Data shown are mean ± SD; n = 5 or 6 chips in each group in **a** and **d**; n = 4 chips in each group in **b** and **c**; unpaired t-test; *** denotes *P* < 0.0001.

Viral infection can trigger alveolar epithelial cell apoptosis.^31^ But macrophages can also induce apoptosis in these cells either directly by expression of TNF-α or TNF-related apoptosis-inducing ligand (TRAIL)^32, 33^ or indirectly by secreting cytokines and chemokines that enhance recruitment of circulatory immune cells that then trigger this programmed death response^34^. When we performed flow cytometry on the cells collected from the apical channel of the chip, we found that the alveolar epithelial cell (CD326^+^CD45^-^) population contained significantly more apoptosis-associated Annexin V/propidium iodide-double-positive (Annexin V^+^/PI^+^) cells in IAV infected chips compared to mock infected chips, both with and without macrophages (**Fig. 2b**). This is consistent with viral infection being the major driver of epithelial cell apoptosis in this model. However, the percent of Annexin V^+^/PI^+^ cells in the chips was significantly greater in chips that contained macrophages (**Fig. 2b**), thus confirming that macrophages also contribute significantly to the apoptotic response. Interestingly, the number of apoptotic cells was significantly lower in mock infected chips when macrophages were present (**Fig. 2b**), indicating that macrophages also play a role in alveolar epithelial cell homeostasis in this *in vitro* model. We then analyzed the macrophage cell (CD326^-^CD45^+^) population after IAV infection and found that there were significantly more Annexin V^+^ (both PI^+^ and PI^-^) cells in infected compared to mock infected chips (**Fig. 2c** and **Supplementary** Fig. 2a, b). This finding suggests that IAV infection induces rapid and massive death of AMs. Apart from alveolar epithelial cell and macrophage cell death, we also observed a significant increase in tissue barrier permeability (**Fig. 2d**) in the alveolus chip in response to viral infection, which was even further increased when macrophages were included (**Fig. 2d**). Taken together, these results suggest that macrophages significantly exacerbate alveolar tissue inflammation and injury by secreting pro-inflammatory cytokines and enhancing alveolar epithelial cell apoptosis during IAV infection.

### Macrophages enhance immune cell recruitment and microvascular injury after influenza infection

Microvascular injury is also a common feature in IAV pathogenesis^3^. Analysis of pro-inflammatory cytokines in the basal effluent from the endothelium-lined vascular channel at 48 h post infection also revealed significant upregulation of IL-6, IL-8, TNF-α, IP-10 and RANTES by IAV infection in chips without macrophages, as well as additional upregulation of IL-1β secretion when macrophages were present (**Fig. 3a**). Flow cytometric analysis of the endothelial cells collected from the basal channel of the chips also showed that IAV infection induced a significant increase in the percent of Annexin V^+^ endothelial cells compared to mock infected chips, both with and without macrophages; however, this apoptotic cell population was higher in infected chips with macrophages (**Fig. 3b**). This was primarily due to the presence of significantly increased Annexin V^+^/PI^-^ cells with a lesser contribution from Annexin V^+^/PI^+^ cells (**Supplementary** Fig. 2c, d).

**Fig. 3.**
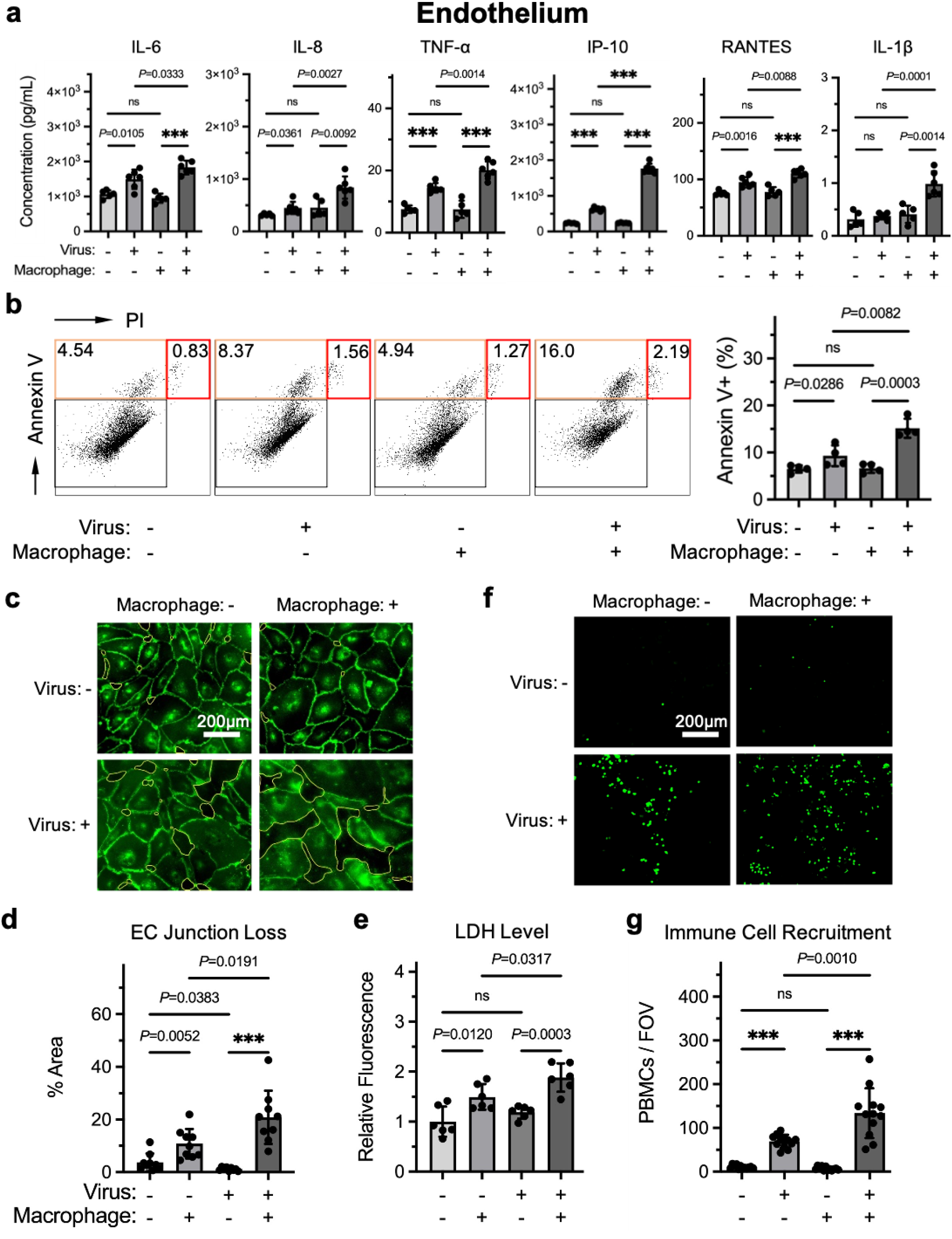
Macrophages exacerbate microvascular inflammation and injury after influenza infection in the alveolus chip. **a** Production of the indicated cytokines at 48 h post infection measured by Luminex assay in the effluent samples collected from infected or mock infected chips, with or without macrophages. **b** Dot plots by flow cytometry analysis (left) for quantifying apoptotic microvascular endothelial cells (Annexin V^+^) (right) in infected or mock infected chips, with or without macrophages, at 48 h post infection. **c** Immunofluorescence with VE-cadherin showing microvascular endothelial cell junction loss as indicated by yellow-outlined area. **d** Image quantification of the area% indicating macrophage-exacerbated microvascular endothelial cell injury after infection. **e** LDH levels measured in the effluent samples collected from infected or mock infected chips, with or without macrophages. **f** Fluorescent microscopic images showing recruitment of CellTracker Green-labeled PBMCs to microvascular endothelial cells in the basal channels. **g** Quantification of the number of recruited PBMCs indicating macrophage-enhanced immune cell recruitment to the alveolar tissue after infection. Data shown are mean ± SD; n = 5 or 6 chips in each group in **a**, **e**; n = 4 chips in each group in **b**; 3 FOVs were analyzed in n = 3 chips in each group in **d**; 4 FOVs were analyzed in n = 3 chips in each group in **g**; unpaired t-test; *** denotes *P* < 0.0001.

In addition, immunofluorescence microscopic studies revealed viral infection induced changes in endothelial cell morphology and loss of cell-cell junctions (**Fig. 3c**) consistent with endothelial cell activation and injury, which were not observed in mock infected chips. Computerized morphometric analysis confirmed that viral infection induced a significant loss of endothelial cell-cell junctions as indicated by an increase in exposed cell-free areas, which was even more increased when macrophages were present (**Fig. 3d**). This is consistent with macrophage-exacerbated microvascular injury and this was supported by detection of significantly increased levels of lactate dehydrogenase (LDH) released into in the basal effluents when macrophages were present (**Fig. 3e**); it is also consistent with the disruption of barrier integrity described above (**Fig. 2d**). In addition, when we performed a peripheral blood mononuclear cell (PBMC) recruitment assay, significantly higher levels of circulating immune cells were recruited to the surface of the pulmonary endothelium in infected compared to mock infected chips, and again, this was further increased when macrophages were present (**Fig. 3f, g**). Together, these results suggest that macrophages significantly increase microvascular injury in response to influenza virus infection and that this may be further enhanced by stimulating recruitment of immune cells to the alveolar tissue.

### Viral infection-induced innate immune responses and macrophage-associated inflammation

We next carried out transcriptomic analyses to further understand the mechanistic pathways underlying macrophage-exacerbated alveolar tissue injury after IAV infection. At 48 h post infection, RNA was extracted from cells in the apical or basal channel in chips with or without macrophages, in the presence or absence of viral infection, and we performed bulk RNA-sequencing (RNA-seq) on the samples. Significant expression of marker genes for type II alveolar epithelial cells (e.g., ABCA3, LAMP3 and SFTPC), type I alveolar epithelial cells (e.g., AGER, HOPX and RTKN2), and AMs (e.g., ITGAX, MRC1 and FCGR1A) was detected in uninfected chips with macrophages (**Supplementary** Fig. 3a), confirming that the Lung Alveolus Chip contains the key cellular components of the human alveolus. In addition, we found significant downregulation of most of these genes following viral infection (**Supplementary** Fig. 3b), suggesting changes in cell differentiation state as well as increased apoptosis after IAV infection, as observed in lungs of patients infected with pandemic SARS-CoV-2^35^.

We next performed a comprehensive analysis of the alveolar epithelial cells and macrophages, where unsupervised principal component analysis (PCA) of RNA-seq clustering data revealed three distinct gene expression clusters (**Supplementary** Fig. 4a). Overall, 5836 differentially expressed genes (DEGs) were identified among 21,219 detected genes in total RNA extracted from the apical channel in infected chips with macrophages compared that from mock infected chips with macrophages, and 6815 DEGs were identified among 21,568 detected genes when comparing RNA from infected chips with macrophages versus those without (**Fig. 4a, b** and **Supplementary** Fig. 4a). Gene set enrichment analysis (GSEA) of KEGG pathways revealed upregulated pathways including cytokine-cytokine receptor interaction, chemokine signaling pathway, and cytosolic DNA sensing pathway, while oxidative phosphorylation and ribosome pathways were downregulated in the alveolar epithelial cells and macrophages after viral infection (**Fig. 4c** and **Table S1**). Similarly, GSEA of hallmark pathways revealed inflammatory response and interferon (IFN)-α response pathways to be upregulated, while oxidative phosphorylation and fatty acid metabolism pathways were downregulated in the alveolar epithelial cells and macrophages after viral infection (**Table S2**). Gene ontology biological process (GOBP) analysis similarly revealed pathways related to antiviral innate immune responses triggered by viral infection, such as upregulated defense response to virus and response to type I IFN (**Supplementary** Fig. 4b and **Table S3**). Notably, our transcriptomic analysis revealed a number of upregulated innate immunity genes in the cells after viral infection, such as MX1, OAS1, TNFSF10, IFI6, CCL5, CXCL10 and IFNB1. This is in line with type I IFN responses and interferon-stimulated gene (ISG) expression as well as pro-inflammatory cytokine production downstream of NF-κB activation in response to signaling mediated by virus-activated retinoic acid-inducible gene-I (RIG-I) and mitochondrial antiviral signaling protein (MAVS)^36^. Taken together, these data show that the human Lung Alveolus Chip faithfully replicates the increased tissue inflammation and altered cell metabolism, as well as induction of antiviral innate immune responses and changes in cell metabolism, that are observed in human patients in response to viral infection of the lung.

**Fig. 4.**
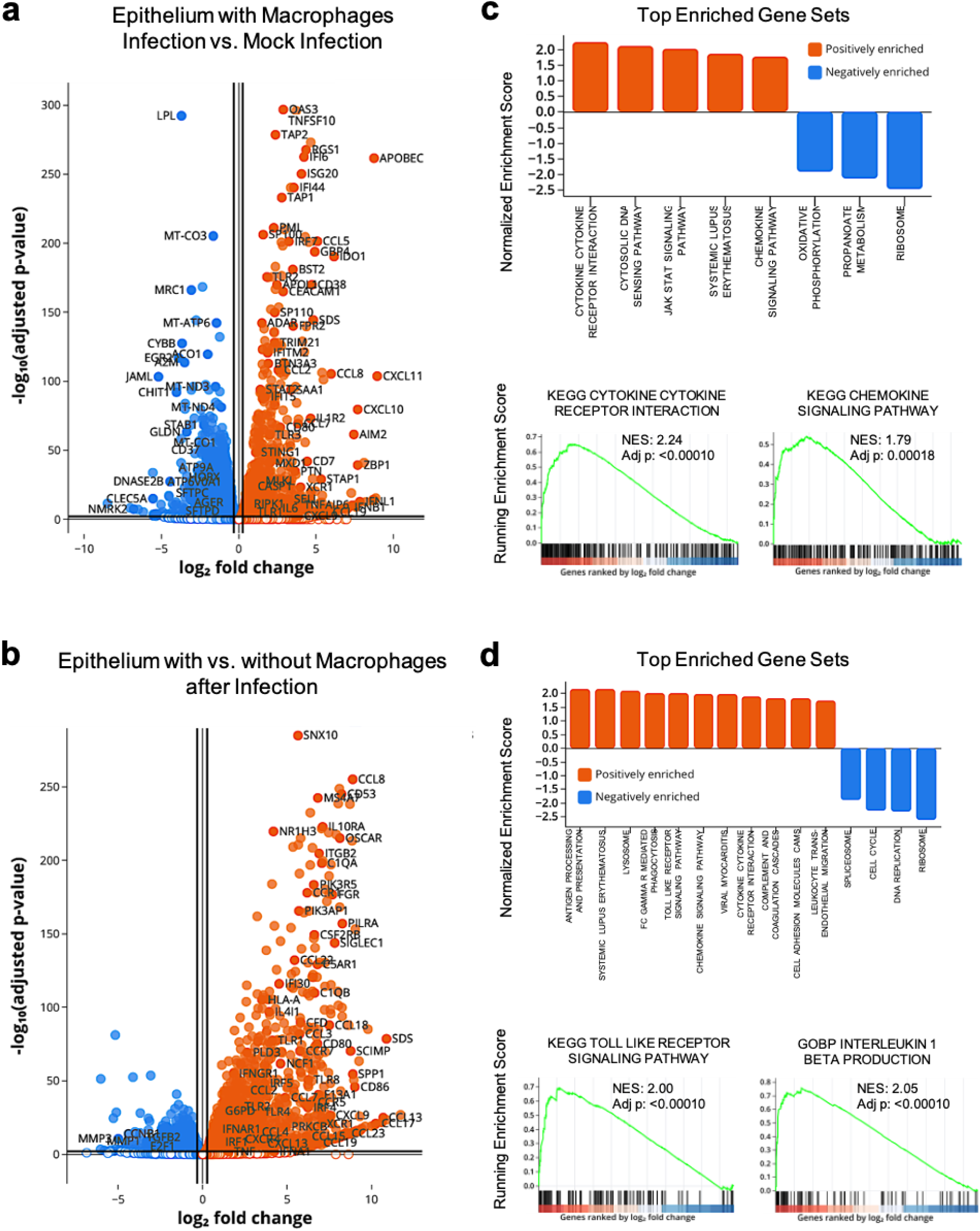
Transcriptomic analyses showing viral infection-induced innate immune responses and macrophage-associated inflammatory pathways in the alveolar tissue. a Volcano plot of DEGs comparing the total RNA extracted from the apical channels in infected chips to that in mock infected chips, both with macrophages. **b** Volcano plot of DEGs comparing the total RNA extracted from the apical channels in infected chips with macrophages to that in infected chips without macrophages. **c** GSEA of KEGG pathways showing viral infection-induced upregulation of cytokine-cytokine receptor interaction, cytosolic DNA sensing pathway, JAK STAT signaling pathway, and chemokine signaling pathway, and downregulation of oxidative phosphorylation, propanoate metabolism, and ribosome in the alveolar tissue. Insets: Enrichment plots are shown for the KEGG cytokine-cytokine receptor interaction and KEGG chemokine signaling pathway gene sets. **d** GSEA of KEGG pathways showing macrophage-associated upregulation of antigen processing and presentation, Fc gamma R-mediated phagocytosis, TLR signaling pathway, chemokine signaling pathway, cytokine-cytokine receptor interaction, complement and coagulation cascades, and leukocyte trans-endothelial migration, and downregulation of cell cycle, DNA replication, and ribosome. Insets: Enrichment plots are shown for the KEGG TLR signaling pathway and GOBP IL-1β production gene sets. n = 4 chips in each group; DEGs were defined with adjusted *P* < 0.01 and fold change < 0.8 or > 1.2; enriched gene sets were defined with adjusted *P* < 0.001; plots were created with Pluto (https://pluto.bio).

To validate the role of macrophages in IAV infection at the transcriptomic level, we analyzed the DEGs in infected chips with and without macrophages (**Fig. 4b**). GSEA of KEGG pathways revealed upregulated pathways including Toll-like receptor (TLR) signaling pathway, chemokine signaling pathway, cytokine-cytokine receptor interaction, leukocyte trans-endothelial migration and antigen processing and presentation, while downregulated pathways included cell cycle, DNA replication and ribosome (**Fig. 4c** and **Table S4**). These results again indicate enhanced alveolar tissue inflammation and immune cell signaling and recruitment, which is consistent with the PBMC recruitment analysis (**Fig. 3f, g**), as well as decreased cell proliferation. Similarly, GSEA of hallmark pathways revealed upregulation of IFN-α response, IFN-γ response and IL2 STAT5 signaling pathways, as well as downregulation of E2F target, G2M checkpoint and mitotic spindle pathways (**Table S5**), again indicating induction of antiviral innate immune responses and cell cycle arrestment. Importantly, GOBP analysis of chips infected with IAV revealed significant upregulation of pathways associated with macrophage-mediated immunopathology, such as myeloid leukocyte activation, myeloid leukocyte mediated immunity, phagocytosis, cell killing and leukocyte mediated cytotoxicity in chips with macrophages compared to those without (**Supplementary** Fig. 4c and **Table S6**). These findings are well aligned with the macrophage-exacerbated lung injury we observed in this model (**Fig. 2b**). GOBP analysis further revealed upregulated TNF superfamily cytokine production and IL-1β production (**Supplementary** Fig. 4c) consistent with the macrophage-enhanced pro-inflammatory cytokine production we observed (**Fig. 2a**).

In addition, we carried out PCA of RNA-seq clustering data from the pulmonary endothelium on-chip, which demonstrated only two distinct gene expression clusters (**Supplementary** Fig. 5a). Overall, 1,857 DEGs were identified among 19,610 detected genes when comparing the endothelial cells in infected versus mock infected chips (both with macrophages). In contrast, only 122 DEGs were identified among 19,218 detected genes when we compared genes in endothelium in infected chips with macrophages versus those without (**Fig. 5a, b** and **Supplementary** Fig. 5a). GSEA of KEGG pathways revealed upregulation of cytosolic DNA sensing and RIG I like receptor signaling pathways, and downregulation of ribosome, lysosome, amino sugar and nucleotide sugar metabolism pathways in endothelium in infected versus mock infected chips (**Fig. 5c** and **Table S7**). Similarly, GSEA of hallmark pathways and GOBP analysis revealed upregulated antiviral defense response as well as downregulated pathways related to altered cell metabolism (**Supplementary** Fig. 5b and **Table S8, S9**). These analyses also identified upregulated antiviral defense response as well as downregulated pathways related to cell cycle arrestment in endothelium in infected chips with macrophages versus those without (**Supplementary** Fig. 5c and **Table S10, S11**). Interestingly, however, GSEA of KEGG pathways comparing the same two groups revealed upregulated complement and coagulation cascades (**Fig. 5d** and **Table S12**). This is important because it suggests that macrophages may contribute to influenza immunopathology by activating the complement system^37, 38^.

**Fig. 5.**
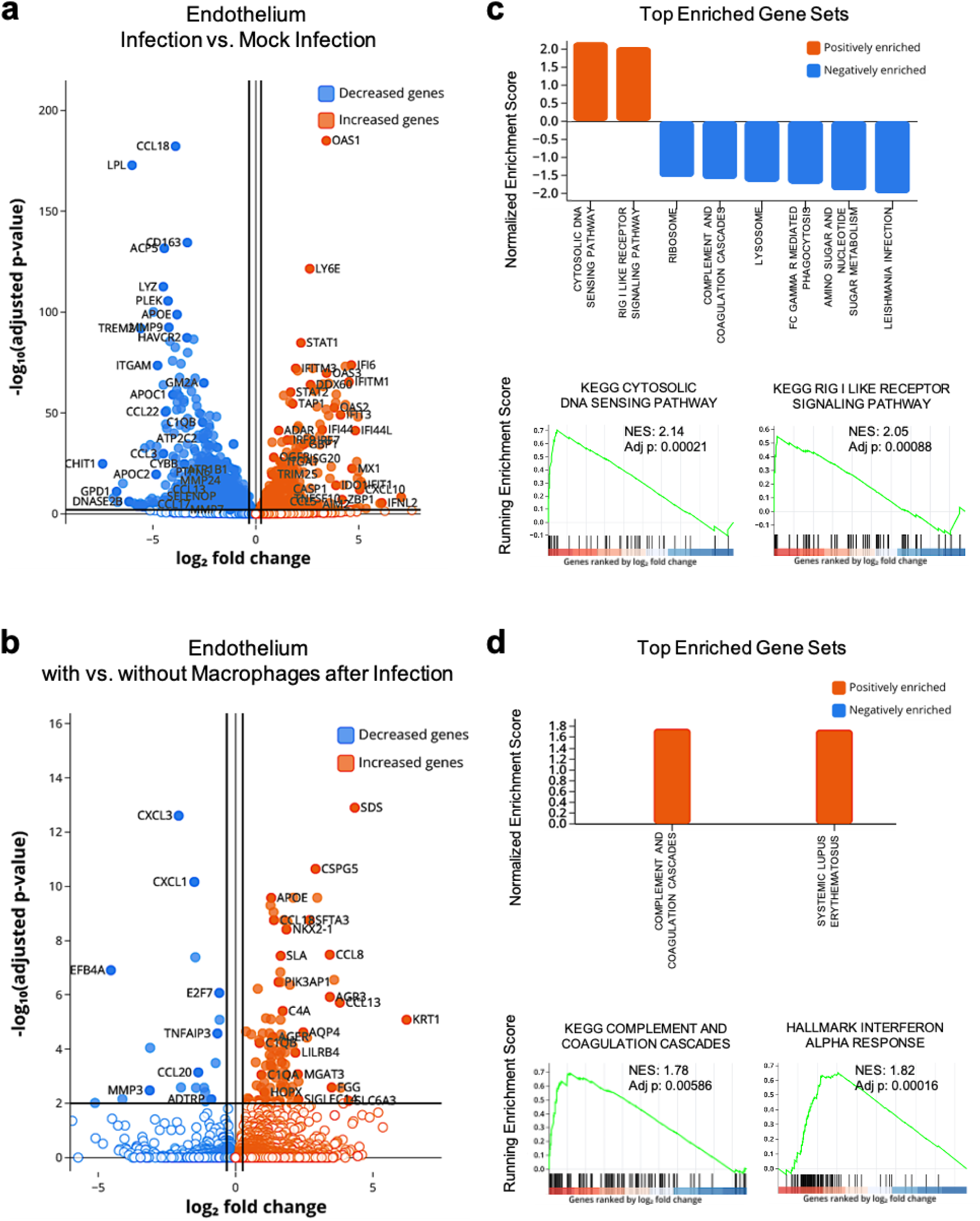
Transcriptomic analyses showing viral infection-induced innate immune responses and macrophage-associated inflammatory pathways in the microvascular tissue. a Volcano plot of DEGs in microvascular endothelial cells in infected chips compared to mock infected chips, both with macrophages. **b** Volcano plot of DEGs in microvascular endothelial cells in infected chips with macrophages compared to infected chips without macrophages. **c** GSEA of KEGG pathways showing viral infection-induced upregulation of cytosolic DNA sensing pathway and RIG I like receptor signaling pathway, and downregulation of ribosome, lysosome, and amino sugar and nucleotide sugar metabolism. Insets: Enrichment plots are shown for the KEGG cytosolic DNA sensing pathway and RIG I like receptor signaling pathway gene sets. **d** GSEA of KEGG pathways showing macrophage-associated upregulation of complement and coagulation cascades, and systemic lupus erythematosus. Insets: Enrichment plots are shown for the KEGG complement and coagulation cascades and hallmark IFN-α response gene sets. n = 4 chips in each group; DEGs were defined with adjusted *P* < 0.01 and fold change < 0.8 or > 1.2; enriched gene sets were defined with adjusted *P* < 0.05; plots were created with Pluto (https://pluto.bio).

### Pyroptosis blockade *via* caspase-1 inhibition prevents influenza virus infection induced lung injury

Our RNA-seq data revealed that mRNA encoding CASP1, the central effector in pyroptosis as well as other DEGs involved in pyroptosis (GO: 0070269), such as ZBP1, AIM2, NINJ1 and GSDMD, are significantly upregulated in samples from the alveolar epithelium in infected Lung Alveolus Chips that contained macrophages compared to non-infected controls (**Supplementary** Fig. 6a, b). Pyroptosis is a highly inflammatory form of lytic programmed cell death that commonly occurs upon infection with intracellular pathogens^39, 40^. RT-qPCR analysis independently confirmed upregulation of CASP1, IL1B, TNF, CXCL10, IL6 and IFNL1 in the RNA extracted from the apical channel (**Supplementary** Fig. 6c), indicating that IAV infection activates both the pyroptosis pathway and lung inflammation on-chip. Moreover, RNA-seq data further detected upregulation of several key genes involved in pyroptosis, such as AIM2, NINJ1, NLRC4, NAIP and PYCARD, in samples from the alveolar epithelium in infected chips that contained macrophages compared to those without (**Supplementary** Fig. 6d). This raised the possibility that blockade of the pyroptosis pathway could potentially prevent macrophage-exacerbated alveolar tissue injury in IAV infection.

We then used the Lung Alveolus Chip as a human preclinical model to explore the therapeutic effect of pyroptosis blockade (**Fig. 6a**). We administered the anti-caspase-1 drug, VX-765 (Belnacasan), which has been approved by the U.S. Food and Drug Administration (FDA) for treatment of human immunodeficiency virus (HIV) infection^41^, Alzheimer’s disease^42^, coronavirus disease 2019 (COVID-19)^37^, atherosclerosis^43^, arthritis^44^ and epilepsy^45^. In initial control studies, we found that when macrophages maintained in submerged cultures in static planar dishes were infected with influenza H3N2 at the same MOI used on chip, a significant increase in caspase-1 protein expression (**Fig. 6b**) as well as NLRP3 speck formation (**Supplementary** Fig. 7a) could be detected by immunofluorescence microscopic imaging at 48 h post infection. Enzyme-linked immunosorbent assay (ELISA) analysis also revealed significant elevation of IL-1β production in the macrophages after IAV infection, which was strongly inhibited by 10 µM VX-765 (**Supplementary** Fig. 7b). VX-765 was then flowed through the basal vascular channel of the Lung Alveolus Chip at the human maximum serum concentration (C_max_, 1.53 µM), mimicking systemic levels of clinical drug treatment. Interestingly, at 48 h post infection, we observed that pyroptosis blockade by VX-765 resulted in increased viral NP RNA levels in the apical wash (**Fig. 6c**), but apoptosis markers in both the alveolar epithelium (Annexin V^+^/PI^+^) and endothelium(Annexin V^+^/PI^+^), as well as LDH levels in the vascular channel, were significantly suppressed (**Fig. 6d, e** and **Supplementary** Fig. 7c), suggesting reduced lung injury.

**Fig. 6.**
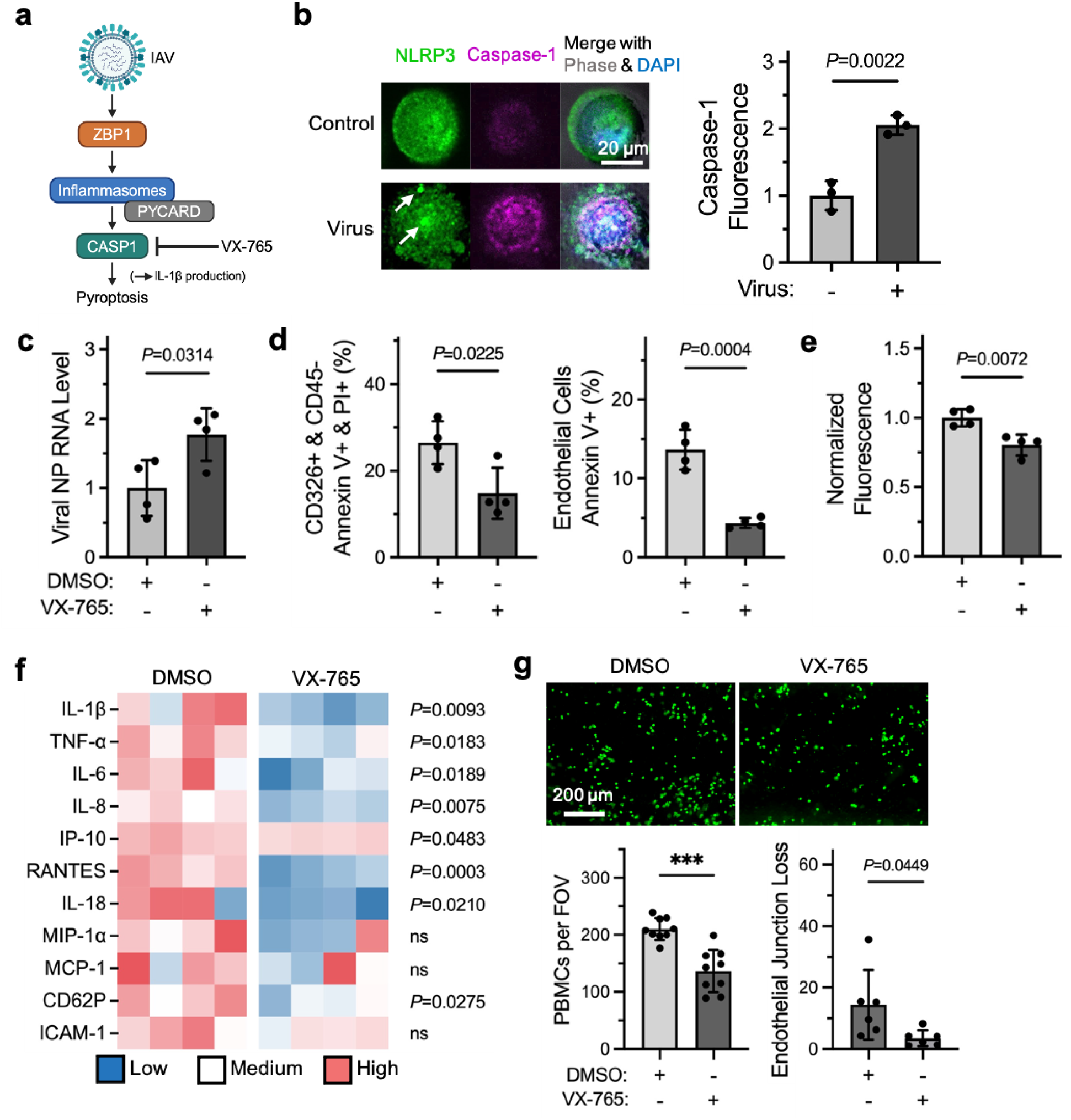
Pyroptotic blockade by caspase-1 inhibition prevents IAV-induced lung injury revealed in the human Lung Alveolus Chip. **a** IAV activates ZBP1 and triggers caspase-1-mediated pyroptosis. Activation of caspase-1 simultaneously leads to IL-1β production. Pyroptosis blockade by caspase-1 inhibitor, VX-765, may present a therapeutic approach for suppressing macrophage-exacerbated lung injury in IAV infection. **b** Fluorescent microscopic images (left) showing NLRP3 speck formation and caspase-1 upregulation in macrophages at 48 h after IAV infection in submerged culture conditions. Image quantification (right) of caspase-1 fluorescence in macrophages. **c** RT-qPCR analysis of relative viral NP RNA levels in the apical wash samples collected from chips treated with VX-765 or DMSO. **d** Graphs showing that VX-765 treatment significantly reduced alveolar epithelial cell (CD326^+^/CD45^-^) and microvascular endothelial cells apoptosis at 48 h after IAV infection. **e** Graphs showing that VX-765 treatment significantly reduced effluent LDH levels at 48 h after IAV infection. **f** Heatmap showing effluent cytokine profiles of chips treated with VX-765 or DMSO at 48 h after IAV infection. **g** Fluorescent microscopic images (top) showing PBMC recruitment to endothelium in chips treated with VX-765 or DMSO at 48 h after IAV infection. Graphs (bottom) showing VX-765 treatment reduced PBMC recruitment and endothelial cell junction loss in the alveolus chip at 48 h after IAV infection. Data shown are mean ± SD; n = 3 wells in each group in **b**; n = 4 chips in each group in **c, d, e** and **f**; 3 FOVs were analyzed in n = 3 chips in each group for PBMC recruitment in **g**; 3 FOVs were analyzed in n = 2 chips in each group for endothelial cell junction loss in **g**; unpaired t-test; *** denotes *P* < 0.0001.

In addition, we analyzed pro-inflammatory cytokine production by the epithelium in the apical wash where we found pyroptosis blockade significantly reduced IL-1β, IL-6, IL-8 and IL-18 levels. A similar analysis of cytokine production by the endothelium in basal channel effluents revealed similar reductions in IL-1β, TNF-α, IL-6, IL-8, IP-10, RANTES, IL-18 and CD62P (**Fig. 6f** and **Supplementary** Fig. 7d). Finally, when we performed PBMC recruitment studies, we found that pyroptosis blockade significantly reduced immune cell recruitment to the endothelium and resulted in less disruption of endothelial cell-cell junctions after IAV infection (**Fig. 6g** and **Supplementary** Fig. 7e). Altogether, these results suggest a strong therapeutic effect of pyroptosis blockade in IAV infection, which ameliorates lung inflammation and injury by reducing pro-inflammatory cytokine production, and immune cell recruitment, as well as apoptosis in both alveolar epithelial cells and endothelial cells.

## Discussion

In this study, we expanded the functionality of a microfluidic human Lung Alveolus Chip by integrating a third cell type, human lung AMs, into the device previously lined by alveolar epithelium interfaced with pulmonary vascular endothelium, which enabled development a more complex and more physiologically relevant model of human distal lung viral infection. Our data demonstrate that the Lung Alveolus Chip faithfully recapitulates host innate immune responses to influenza H3N2 infection at the morphological, biochemical, and transcriptomic levels. Moreover, our transcriptomic analyses are in agreement with our findings at the protein level and cellular level, where macrophages significantly exacerbated lung inflammation and injury by secreting pro-inflammatory cytokines (e.g., IL-1β and TNF-α), promoting alveolar epithelial cell and endothelial apoptosis as well as enhancing immune cell recruitment in response to IAV infection. This model also allowed us to demonstrate that while the presence of AMs helps to reduce viral load, it also exacerbates lung injury in response to IAV infection. This effect appears to be mediated by virus-triggered pyroptosis of AMs, which causes release of pro-inflammatory mediators, elevates epithelial and endothelial cell apoptosis, disrupts the pulmonary tissue barrier, and enhances infiltration of immune cells. The rapid and massive macrophage deaths induced by IAV infection were associated with inflammasome assembly activation and caspase-1 upregulation that have been shown to induce pyroptosis and IL-1β release, which in turn amplifies host inflammatory responses and enhances viral clearance^46^. In contrast, when macrophages are absent, levels of IL-1β and other pro-inflammatory mediators are lower, but viral titers are higher. Importantly, our data further revealed that pharmaceutical inhibition of caspase-1, a critical pyroptotic pathway mediator, suppresses lung inflammation and injury after IAV infection. This finding suggests that pyroptosis blockade may represent a potential therapeutic approach that can be used alone or in combination with other antiviral and/or anti-inflammation drugs to treat life-threatening lung illnesses caused by IAV infection. Interestingly, SARS-CoV-2 infection also triggers pyroptosis in monocytes and lung-resident macrophages, but not in infected epithelial cells or endothelial cells^37, 47^, and this causes systemic inflammation that is a major contributor to COVID-19 pathogenesis^48^. Thus, pyroptosis blockade might also be of great therapeutic value for suppressing hyper-inflammatory host responses associated with other respiratory viruses, including SARS-CoV-2.

AMs are the major sentinel cells in the alveolar space that critically elicit lung inflammation; however, how they crosstalk to other tissue cells and immune cells in the lung alveolus is incompletely understood. Both our cytokine and RNA-seq analysis revealed that IL-1β and TNF-α production are upregulated in Lung Alveolus Chips infected with IAV when macrophages are present, Thus, these sentinel cells appear to communicate to alveolar epithelium, microvascular endothelium and circulatory immune cells by producing IL-1β and TNF-α. This is consistent with the finding that high levels of IL-1β and TNF-α are found in lung specimens of critically ill patients with severe influenza and COVID-19^49^. On one hand, IL-1β is known to upregulate adhesion molecules on endothelial cells which has multiple effects, including increasing vascular permeability^50^; stimulating pro-inflammatory cytokine production in monocytes, macrophages, dendritic cells, mast cells and basophils^34^; promoting neutrophil survival and protease release^51^; and stimulating IFN-γ production by T cells as well as T helper 17 cell differentiation^52^. On the other hand, TNF binding to its receptors activates the NF-κB pathway, which induces potent cellular inflammatory responses in a number of immune and non-immune cells^53^, as well as caspase-8-mediated apoptosis and RIPK3-mediated necroptosis^54^. These past findings may help to explain why we observed significantly more apoptosis in the alveolar epithelium and endothelium, along with enhanced tissue barrier disruption and immune cell recruitment when macrophages were present in IAV infected chips.

One of the most clinically relevant findings of this study was that inhibition caspase-1, which suppresses IL-1β production and pyroptotic cell death, prevented lung injury in our human model of IAV infection. However, we found that viral replication in the alveolar space increased following pyroptosis blockade on-chip, which is potentially due to the fact that pyroptosis and the released pro-inflammatory mediators play key roles in innate immune mechanisms for viral clearance. These results highlight the importance of the timing of drug dosing, which needs to be carefully evaluated to avoid inhibiting viral clearance during early stages of lung infection. However, it is important to note that administration of VX-765, the same caspase-1 inhibitor tested in this study, starting as early as day 2 post infection, significantly suppressed immune activation and reduced plasma viral load in a mouse model of HIV-1 infection^41^. Thus, targeting of caspase-1 for pyroptosis blockade early after infection, either alone or in combination with antiviral drugs, may represent a viable path for treatment of viral respiratory diseases and preventing pneumonia and lung injury.

The primary focus of this study was to integrate AMs, a previously missing cell component of human alveoli, into the Lung Alveolus Chip so that the host innate immune responses against influenza infection can be more accurately modeled *in vitro*. Because it is impossible to experiment on humans, preclinical drug studies are often conducted on animals. However, in the context of respiratory viral infection, mouse models do not reflect the pathophysiology of the natural human host. Many of the respiratory viruses that infect humans and cause ARDS, such as IAV and SARS-CoV-2, do not naturally infect mice. Moreover, mouse studies typically employ survival as the primary endpoint, but not severity of pneumonitis which is more human-relevant. Nonhuman primate models provide a more beneficial benchmark for investigating human respiratory viral infections^55^; however, their use is limited by short supply, long breeding periods, high costs, and serious ethical concerns. Thus, there is a great need for human preclinical models that recreate the human lung alveolar-capillary interface with ALI and residential immune cells. Earlier studies have attempted to model diseases of the human alveolus with AMs present *in vitro*^56, 57, 58^; however, they either utilized cancerous cell lines or employed static models that lack dynamic fluid flow, which is a key feature of human lung physiology. As a result, these past models did not accurately recapitulate physiologically relevant host inflammatory responses or the complex interplay between different cell types. The Lung Alveolus Chip model described here that includes human AMs overcomes these limitations, thus providing a useful preclinical model for gaining greater insights into cellular mechanisms and crosstalk underlying viral infection-induced lung injury. Given the recently approved FDA Modernization Act 2.0 that permits the FDA to consider data generated in human Organ Chips in future investigational new drug (IND) applications, this model is also well suited for rapid identification and optimization of new and more effective therapeutics for preventing development of viral pneumonia and ARDS.

It should be noted that this study has certain limitations. First, we used macrophages that were differentiated from marrow-derived monocytes by addition of M-CSF and IL-4. Although in certain lung conditions, AMs can originate from marrow-derived monocytes, their surface markers and functions are different due to the impact of local lung environment^59^. However, a recent study uncovered that M2 macrophages bioenergetically resemble macrophages found in bronchial alveolar lung fluid (BALF) in healthy volunteers^60^, which supports the use of polarized macrophages as a model to study AM function *in vitro*. Moreover, the modular feature of the human Organ Chip system could allow primary AMs obtained in BALF samples collected from healthy or diseased individuals to be integrated into the chips to compare the outcome of viral infection in future studies. This could enable analysis of the impact of aging on lung infections as monocytic macrophages are dominant in aged lungs. Second, the presence of circulatory immune cells during the entire IAV infection process is missing in the present model as we only introduced PBMCs 48 h post infection as an endpoint recruitment assay. Numerous studies have demonstrated that innate immune cells such as neutrophils and monocytes are key contributors to influenza lethality, and their infiltration and activation are closely related to pyroptosis pathways^61, 62^. Thus, future studies should be carried out to uncover the contribution of AM-orchestrated recruitment of inflammatory neutrophils and monocytes and to study the underlying control mechanisms, which could lead to identification of potential new therapeutic targets. Finally, our studies were carried out in the absence of breathing motions to focus on the specific roles of macrophages independent of known effects of mechanical forces on suppressing macrophage inflammation^24^. The contribution of physiological breathing motions, as well as abnormal breathing motions (e.g., that might occur in an infected patient in the intensive care unit), should be explored in future studies to obtain an even higher-level understanding of all the processes that are involved in the human host response to lung infection.

## Methods

### Human Lung Alveolus Chip

Two-channel microfluidic Organ Chip devices were purchased from Emulate Inc (Boston, MA). On day 0, chips were activated using ER1 and ER2 reagents (Emulate) following manufacturer’s instruction and both channels were coated with 200 μg/ml collagen IV (5022-5MG, Advanced Biomatrix, Carlsbad, CA) and 15 μg/ml laminin (L4544-100UL, Sigma Aldrich, St. Louis, MO) at 37°C overnight. On day 1, chips were washed with EGM-2MV medium (CC-3202, Lonza, Basel, Switzerland), and human primary lung microvascular endothelial cells (CC-2527, Lonza; obtained at passage 3 and expanded according to passage 5 before use) at 8 million/mL were seeded into the basal channel. Chips were then flipped to allow EC adhesion to the basal side of the porous membrane. Following 1 h incubation at 37°C, chips were flipped back and human primary lung alveolar epithelial cells (H-6053, Lot#F040615Y72NX, Cell Biologics, Chicago, IL; obtained at passage 3 and used immediately after thawing without further expansion) at 0.5 million/mL were seeded into the apical channel. Following 1 h incubation at 37°C allowing alveolar epithelial cell adhesion to the apical side of the membrane, basal channels were washed with EGM-2MV and apical channels were washed with human epithelial cell growth medium (H-6621, Cell Biologics) for removing non-adhered cells. Chips were maintained under static condition at 37°C overnight. On day 2, chips were inserted into the Pods (Emulate Inc.) and connected to the Zoë system to ensure perfusion. Apical channels and basal channels were perfused with human epithelial cell growth medium and EGM2-MV, respectively at a flow rate of 30 μL/h. On day 5, 1 μM dexamethasone was added to the medium in apical channels to enhance epithelial barrier function. On day 7, medium in apical channels was removed by flowing at 1000 μL/h for 2 min for establishment of ALI. From this timepoint, chips were only fed through the basal channel, and the basal medium was switched to EGM-2MV with 0.5% fetal bovine serum (FBS). 4 days later, chips were used for infection experiments. In experiments included in **Figs. 4-6**, the alveolar epithelial cells were isolated from an alternative donor source as previously described^63^. The donor tissue was obtained from the International Institute for the Advancement of Medicine (IIAM) and used in accordance with the regulations by IIAM. The alveolar epithelial cells isolated from the donor were directly used without expansion and were introduced into apical channels at 5 million/mL. Cells were maintained in K+DCI medium as previously described^64^ before switching to ALI culture. All uses of human materials have been approved by Harvard Longwood Campus Institutional Review Board (IRB).

### PBMC isolation and macrophage differentiation

De-identified human patient-derived apheresis collars (by-product of platelet isolation) were obtained from the Crimson Biomaterials Collection Core Facility under approval IRB protocols (#22470). Informed written consent was not required. Human PBMCs were isolated by density centrifugation using Lymphoprep (07801, StemCell Technologies, Vancouver, Canada). Some of the isolated PBMCs were subsequently cryopreserved in a solution of 90% FBS and 10% DMSO for PBMC recruitment assay. CD14^+^ human monocytes were isolated from the PBMCs using an EasySep™ Human Monocyte Enrichment Kit (19058, StemCell Technologies). The isolated monocytes were cultured in RPMI medium supplemented with 10% FBS and 1% Antibiotic-Antimycotic (A-A, 15240062, Thermo Fisher Scientific, Waltham, MA) in T75 flasks at 2 million/mL and differentiated for 7-9 days with 50 ng/mL recombinant human M-CSF (300-25, PeproTech, Cranbury, NJ) and 20 ng/mL recombinant human IL-4 (200-04, PeproTech). Following differentiation, the human macrophages were incubated with 10 mM ethylenediaminetetraacetic acid (EDTA, 15575020, Thermo Fisher) for 30 min at 37°C, after which the cells were harvested using cell scrapers. Some of the macrophages were seeded at 1 million/mL into the apical channel and incubated statically for 2 h at 37°C prior to infection experiments, and the rest were cryopreserved for separate analysis.

### Viruses

One pandemic influenza strain A/Hong Kong/8/68 (H3N2) was employed in this study, which was purchased from ATCC (VR-1679, Manassas, Virginia). Viruses were propagated in MDCK cells (CCL-34, ATCC) following established protocols^65^. Viral titers of the stock solution and chip samples were determined by plaque assays.

### Plaque assay

Plaque assays were performed following established protocols^66^. Briefly, confluent monolayers of MDCK cells in 12-well plates were washed with DPBS without calcium or magnesium (14190250, Thermo Fisher) and the MDCK infection medium (DMEM supplemented with 1% A-A and 1X non-essential amino acids) and inoculated with 0.2 mL of 10-fold serial dilutions of influenza virus samples for 1 h at 37°C. The inoculums were then removed and the cells were washed with the MDCK infection medium. Meanwhile, a solution was prepared by mixing 3.9 mL of 2% agarose (A9539, MilliporeSigma, Burlington, MA), 9.1 mL of the overlay medium (10X MEM diluted with sterile water and supplemented with 7.5% bovine serum albumin (BSA) fraction V, 200 mM L-glutamine, 7.5% sodium bicarbonate, 1% DEAE-dextran and 1% A-A), and 2 µg/mL of TPCK-treated trypsin, which were thereafter overlaid on top of the MDCK cells with 1 mL per well. The plates were then incubated at room temperature (RT) for 10 min to allow the agarose gel to cool and solidify, after which the plates were flipped and incubated at 37°C to enable plaque formation. 3 days later, the cells were fixed with 4% paraformaldehyde (PFA, Electron Microscopy Sciences, Hatfield, PA) for 30 min at RT, and the agarose gel was carefully removed by vacuum. Finally, fixed cells were washed with DPBS and stained with crystal violet (470300-938, VWR, Radnor, PA) to visualize the plaques. Viral titers were determined as plaque-forming units per milliliter (PFU/mL).

### Chip infection experiments

Chips and the Pods were removed from the Zoë system, and chips were disconnected from the Pods in a biosafety cabinet. 35 μl of viral inoculum was introduced into the apical channel to infect the alveolar epithelial cells with or without macrophages. Chips were then incubated statically at 37°C. After 1-h incubation, chips were removed from the Pods and apical channels were emptied. Thereafter, chips were re-plugged into the Pods, re-connected to the Zoë system, and resumed to flow culture at 30 μL/h under ALI.

### Immunofluorescence and confocal microscopy

Cells were washed with DPBS, fixed with 4% PFA for 15 min at RT, permeabilized with 0.1% Triton X-100 (X100, Sigma Aldrich) for 15 min, and blocked with 10% goat serum (50062Z, Thermo Fisher) in DPBS for 1 h at RT. Cells were then incubated with primary antibodies in DPBS (1:100 dilution) with 5% goat serum and 0.1% Triton X-100 overnight at 4°C, followed by incubation with secondary antibody in DPBS (1:500 dilution) with 5% goat serum and 0.1% Triton X-100 for 1 h at RT. If using conjugated antibodies for immunofluorescence, cells were incubated with the antibodies in DPBS (1:50 dilution) with 5% goat serum and 0.1% Triton X-100 for 1 h at RT overnight at 4°C. Cell nuclei were stained with 1 μg/mL 4′,6-diamidino-2-phenylindole (DAPI, D9542, Sigma Aldrich) for 10 min at RT before imaging. Imaging was conducted using a confocal laser-scanning microscope (Leica SP5 X MP DMI-6000, Wetzlar, Germany) and image processing was done using the ImageJ software (v 2.14.0). Antibodies used in this study can be found in Table S13.

### Apical wash and RNA extraction

To collect apical wash, a fresh, empty 200 μL filtered pipette tip was inserted into the outlet pore of the apical channel, and 100 μL of DPBS was introduced into the apical channel through the inlet pore using another 200 μL filtered pipette tip. The apical channel was washed once with the DPBS, which was collected back to the filtered pipette tip at the inlet. The apical wash sample was finally pipetted into a fresh Eppendorf tube and stored at -80°C. To lyse the epithelial cells on chip, a fresh, empty 200 μL filtered pipette tip was inserted into the outlet pore of the apical channel, and 100 μL Buffer RLT Plus (1053393, Qiagen, Hilden, Germany) was introduced into the apical channel through the inlet pore using another 200 μL filtered pipette tip. The micropipette plunger was quickly pressed and released for five times, after which the cell lysates were collected into a fresh Eppendorf tube and stored at -80°C. Endothelial cell lysates were subsequently collected from the basal channel similarly and stored at -80°C. RNA was extracted from the cell lysates using RNeasy® Plus Micro Kit (74034, Qiagen) and stored at - 80°C.

### RT-qPCR

After determining RNA concentrations by spectrophotometry, 500 ng of total RNA was used for cDNA synthesis. Reverse transcription was conducted using the Omniscript RT Kit (205113, Qiagen). Quantitative real-time PCR was performed using the SsoAdvanced Universal SYBR Green Supermix (1725272, Bio-Rad Laboratories, Hercules, CA) on the Quantstudio 7 Flex Real-Time PCR system (4485701, Thermo Fisher). The specificity of primers was confirmed by melting curve analysis. Relative RNA level was quantified using the ΔΔCt method^67^ and normalized to the endogenous control GAPDH. All primers were purchased from IDT (Newark, NJ) as shown in Table S14.

### Cytokine analysis

Chip outlet reservoirs were emptied before infection experiments and chip basal effluents were collected at 48 h post infection. Cytokine levels in the effluents were analyzed using custom ProcartaPlex assay kits (Thermo Fisher). The analyte concentrations are evaluated using a Bio-Plex 3D suspension array system and analyzed for standard curve fitting and concentration calculations with Bio-Plex Manager software (Bio-Rad, v 6.0).

### Flow cytometry

Cells in chips were washed with DPBS and were incubated with ACCUMAX cell detachment solution (07921, StemCell Technologies) for 30 min at 37°C. Cells were then collected in DMEM with 10% FBS and 1% A-A, washed with cold DPBS, and re-suspended in 100 µL cold stain buffer (554656, BD Biosciences, Franklin Lakes, NJ) with 1% FcR blocking reagent (Miltenyi Biotec, Gaithersburg, MD) on ice. 1 µL of fluorescent conjugated antibodies against human CD326 and CD45 were added to the suspension of cells from the apical channel, and were incubated for 30 min on ice. Cells from the basal channel remained unstained during this step. All cells were then washed once with cold stain buffer and washed twice with cold annexin V binding buffer (V13246, Thermo Fisher). Thereafter, cells were re-suspended in 100 µL cold annexin V binding buffer, and 5 µL of FITC annexin V (640906, BioLegend, San Diego, CA) and 1 µL of 1 mg/mL propidium iodide (P1304MP, Thermo Fisher) in annexin V binding buffer were added into the cell suspension. Following 15 min RT incubation, 400 µL of annexin V binding buffer was added to the cell suspension, and cells were kept on ice before analyzed using CytoFlex LX (Beckman Coulter, Brea, CA). Data were analyzed using FlowJo V10 software (Flowjo, LLC, Ashland, OR).

### LDH assay

LDH levels in the chip basal effluents collected at 48 h post infection was analyzed as a secondary measurement of cell deaths using CytoTox 96® Non-Radioactive Cytotoxicity Assay (G1780, Promega, Madison, WI) following manufacturer’s instruction.

### Barrier function analysis

In some experiments, barrier function was analyzed in chips at 48 h post infection. All reservoirs were emptied before carrying out the assay, and fresh, warm medium (EGM-2MV with 0.5% FBS) was introduced into both channels in the chip and also into the apical channel inlet reservoir. Dextran, Cascade Blue (D7132, Thermo Fisher) was diluted in fresh, warm medium to a final concentration of 100 μg/mL, and 1 mL of the dosing medium was added into the basal channel inlet reservoir. Thereafter, both channels were flushed at 500 μL/h for 10 min at 37°C to allow the dosing medium to fill the basal channel. Outflows were then removed from the outlet reservoirs, and the chips were flowed at 120 μL/h for both channels for 2 h at 37°C. Outflows were collected from the outlet reservoirs and the fluorescence was analyzed at 380/420 (excitation/emission). The apparent permeability of the alveolar-capillary tissue barrier was calculated using the formula: Papp = J / (A × ΔC), where Papp is the apparent permeability, J is the molecular flux, A is the total area of diffusion and ΔC is the average gradient.

### PBMC recruitment assay

PBMCs isolated as stated above were thawed and resuspended in 10 mL RPMI medium supplemented with 1% A-A. 10 μL CellTracker Green (C7025, Thermo Fisher) at a final concentration of 10 μM was used to label the PBMCs for 20 min at 37°C. Cells were then centrifuged, washed with fresh RPMI medium supplemented with 10% FBS and 1% A-A, and resuspend in EGM-2MV with 0.5% FBS at 20 million/mL. At 48 h after infection, chips were detached from Pods and 25 μl of the PBMC suspension was injected into the basal channels. Chips were thereafter placed on chip cradles and flipped. PBMCs were allowed to adhere at 37°C for 2 h, after which the chips were flipped back and washed with 200 μl fresh suspension medium to wash out unattached cells. Chips were immediately imaged under a fluorescent microscope.

### Bulk RNA sequencing and bioinformatic analysis

RNA-seq was performed by Azenta Life Sciences (Burlington, MA) using an RNA-seq package including polyA selection and sequencing using an Illumina HiSeq for 150 bp pair-ended reads. These sequence reads were trimmed using Trimmomatic v.0.36 to eliminate adapter sequences and nucleotides with poor quality. Subsequently, these trimmed reads were mapped onto the Homo sapiens GRCh38 reference genome using STAR aligner v.2.5.2b. Unique gene hit counts were computed using Counts from the Subread package v.1.5.2. Genes were filtered to include only genes with at least 3 reads counted in at least 20% of samples in any group, followed by differential expression analysis using DESeq2 R package, which tests for differential expression based on a model using the negative binomial distribution. The false discovery rate (FDR) method was applied for multiple testing correction. DEGs were identified if the gene’s FDR-adjusted *P*-value is < 0.01 and its fold change is either less than 0.8 or greater than 1.2. GSEA was performed using the fgsea R package and the fgseaMultilevel() function. Gene set collection, including Hallmarks, Canonical pathways - KEGG, and Gene Ontology gene sets - biological process, from the Molecular Signatures Database (MSigDB) was curated using the msigdbr R package4. Prior to running GSEA, the list of gene sets was filtered to include only gene sets with between 5 and 1000 genes. CPM values were calculated by dividing the number of reads mapped to a gene by a million-scaling factor divided by the total of mapped reads, using the cpm() function in the edgeR R package with log=F. Analysis performed using Pluto software (https://pluto.bio).

### Drug study

VX-765 (S2228, Selleck Chemicals LLC, Houston, TX) was purchased. Blood C_max_ of VX-765 was obtained from Sigma Aldrich (https://www.sigmaaldrich.com/US/en/product/mm/531372). VX-765 was diluted in EGM-2MV with 0.5% FBS at a final concentration of 1.53 µM and dosed through the basal channel at the time point indicated in **Fig. 6c**. Equal amount of DMSO was diluted in the medium and used in the control group. Final DMSO concentration in each group is less than 0.1%.

### Statistics

All reported experiments (excluding RNA-seq) were performed at least twice, with at least three technical repeats in each group in each experiment. Data is represented as mean values ± standard deviation (SD). Graphing and statistical comparison of the data were performed using GraphPad Prism (v 10.1.0). Statistical significance for two-group comparisons were determined using unpaired t-test. *P* < 0.05 were considered to be statistically significant; exact *P* values were labeled in all figures.

## Data availability

All bulk RNA-seq data have been uploaded to the National Center for Biotechnology Information (NCBI) database and made publicly available with the accession number GSE272710. Additional data are available in the Supplementary Information. The data were analyzed using available data packages mentioned in the manuscript and no new or custom codes were created to analyze the data. There are no restrictions on data availability and raw data has been made available. Source data are provided within this manuscript.

## Acknowledgements

This research was supported by Defense Advanced Research Projects Agency (DARPA) under Cooperative Agreement (HR0011-22-2-0017) awarded to D.E.I, National Institutes of Health (NIH) 3UH3HL141797-04S1 awarded to D.E.I, and Wyss Institute for Biologically Inspired Engineering at Harvard University. The authors thank Maurice Perez, Eric Zigon and Thomas C. Ferrante for lab support and instrument training, Rani K. Powers and Megan M. Sperry for discussion on RNA-seq data analysis and Pluto software, and Keleigh M. Quinn and Kenneth Carlson for grant and project management.

## Author contributions

Y.M., Y.Z., H.B., S.H. and D.E.I conceived the study. G.G., C.B., and S.H. oversaw the study. Y.M., Y.Z., A.J., H.B., A.G., G.G., C.B., S.H. and D.E.I. participated in experiment design and/or discussions. Y.M., Y.Z., A.J., G.E.M. and C.B. performed the experiments and data collection. Y.M., R.P. and R.J.M acquired confocal microscopy images. H.B. performed alveolar cell isolation from the donor tissue obtained from IIAM. Y.M. and Y.Z. analyzed the data, prepared the figures and tables, and wrote the manuscript with input from all authors. D.E.I. reviewed and edited the manuscript. All authors approved the final version.

