## Supplement Information for "Exacerbation of influenza virus induced lung injury by alveolar macrophages and its suppression by pyroptosis blockade in a human lung alveolus chip"

[4]Equal contribution

**Supplementary Figures**

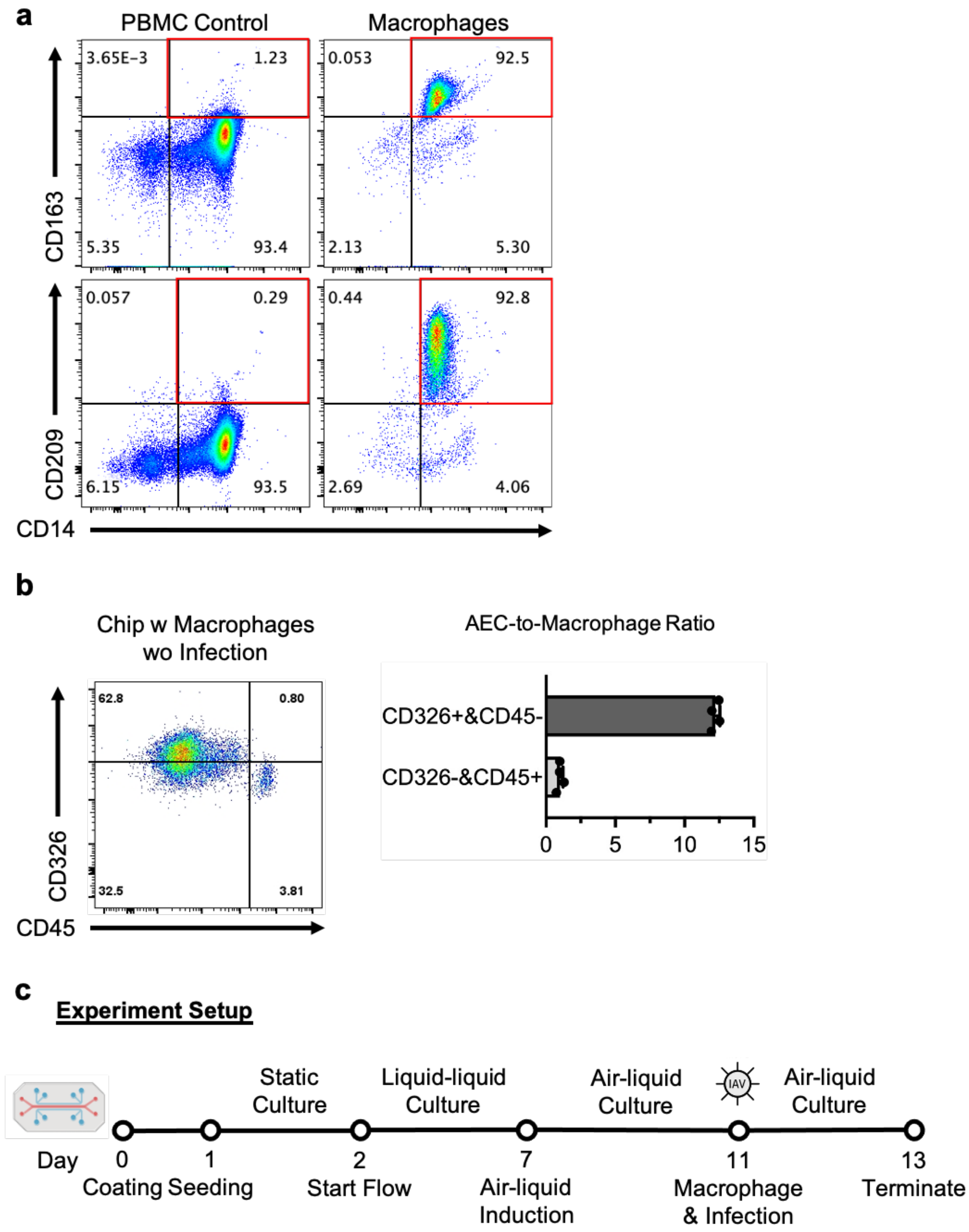

Figure S1. (a) Dot plots by flow cytometry analysis showing differentiated cells expressing macrophage markers, CD163 and CD209. PBMCs were used as control samples. (b) Dot plot (left) by flow cytometry analysis showing AECs ($CD326^+/CD45^-$) and macrophages ($CD326^-/CD45^+$). Graph (right) showing analysis of AEC-to-macrophage ratio on chip. (c) Illustration of experimental protocol for culturing the alveolus chip and its infection.

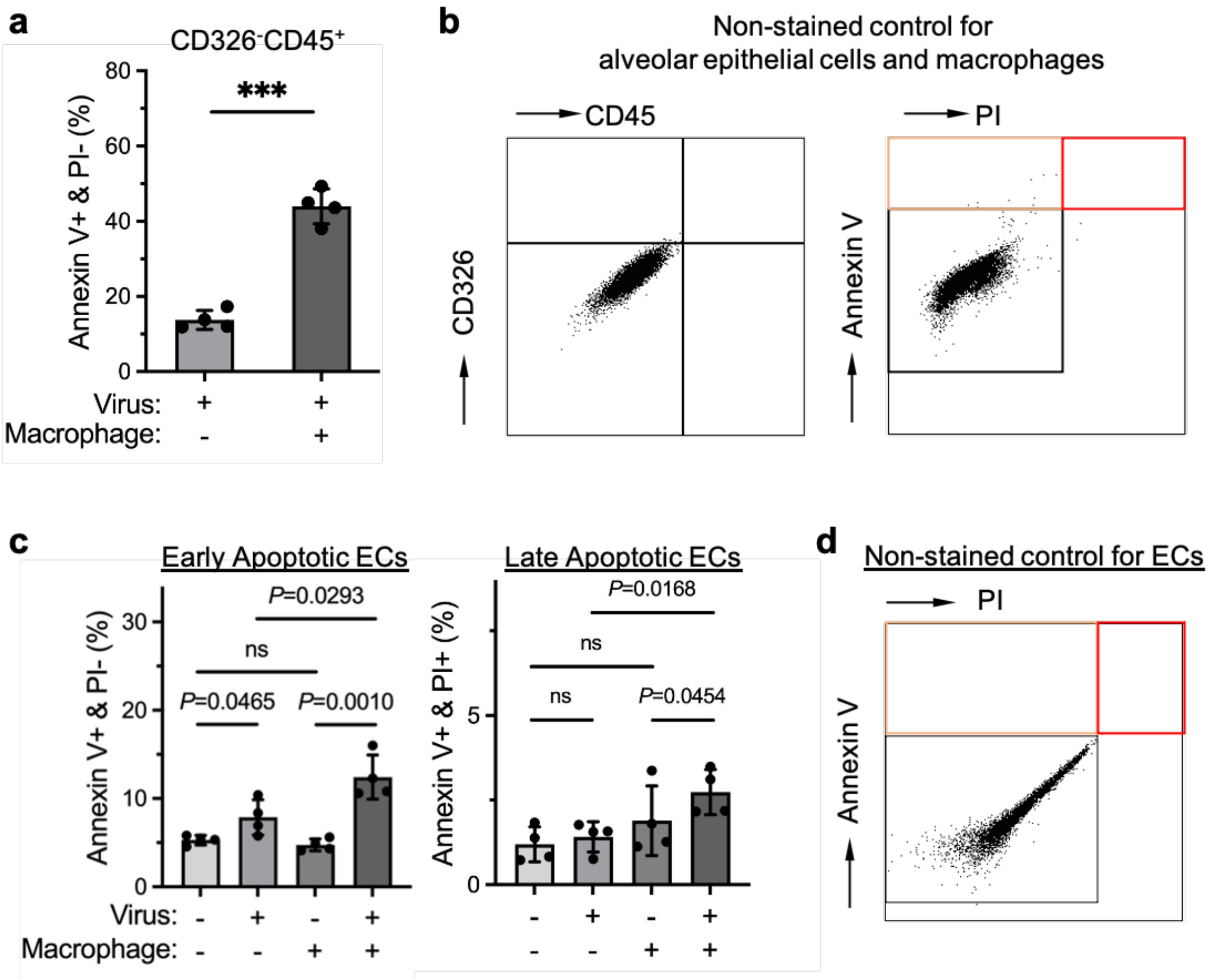


Figure S2. (a) Graph showing early apoptotic macrophage (Annexin V$^+$/PI$^-$) in infected or mock infected chips, at 48 h post infection. (d) Dot plots by flow cytometry analysis of a non-stained control sample from the apical channel of a mock infected chip with macrophages. (c) Graphs showing early apoptotic ECs (Annexin V$^+$/PI$^-$) and late apoptotic ECs (Annexin V$^+$/PI$^+$) in infected or mock infected chips, with or without macrophages, at 48 h post infection. (d) Dot plot by flow cytometry analysis of a non-stained control sample from the basal channel of a mock infected chip with macrophages. Data shown are mean ± SD, n = 4 chips in each group in (a) and (c); unpaired t test; *** denotes $P < 0.0001$.

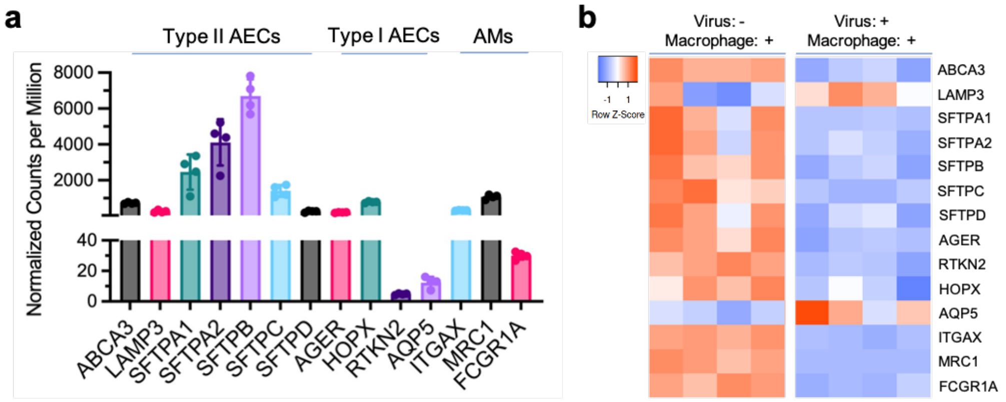


Figure S3. (a) Type II AEC, type I AEC and AM marker gene expression in mock infected chips with macrophages. (b) Heatmap showing IAV infection-induced downregulation of type II AEC marker genes, including ABCA3 and SFTPC, type I AEC marker genes, including AGER and RTKN2, and AM marker genes, including ITGAX and MRC1, in the alveolus chip with macrophages. n = 4 chips in each group in the RNA-seq analysis.

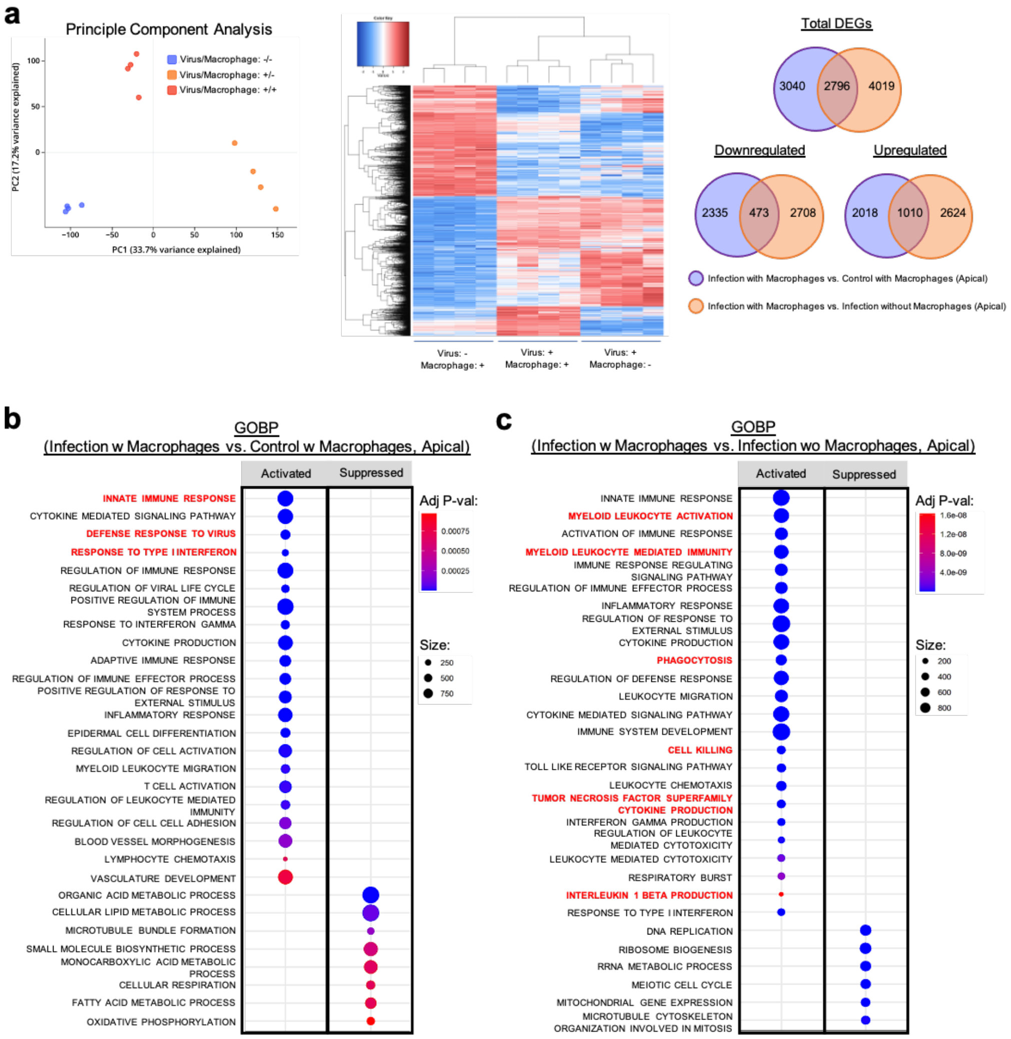


Figure S4. (a) Plot (left) showing principal component analysis of the alveolar tissue with different clusters formed by the three indicated groups. Heatmap (middle) showing clustering analysis of DEGs among the three groups. Venn diagrams (right) depicting the shared or unique DEGs between each comparison. (b) Enriched gene sets revealed by GOBP analysis of DEGs in the alveolar tissue in infected chips compared to mock infected chips, both with macrophages. 30 out of 118 are shown. (c) Enriched gene sets revealed by GOBP analysis of DEGs in the alveolar tissue in infected chips with macrophages compared to infected chips without macrophages. 30 out of 411 are shown. The color of the dots represents the significance of enrichment and the size of the dots represents the input number of each term in (b) and (c).

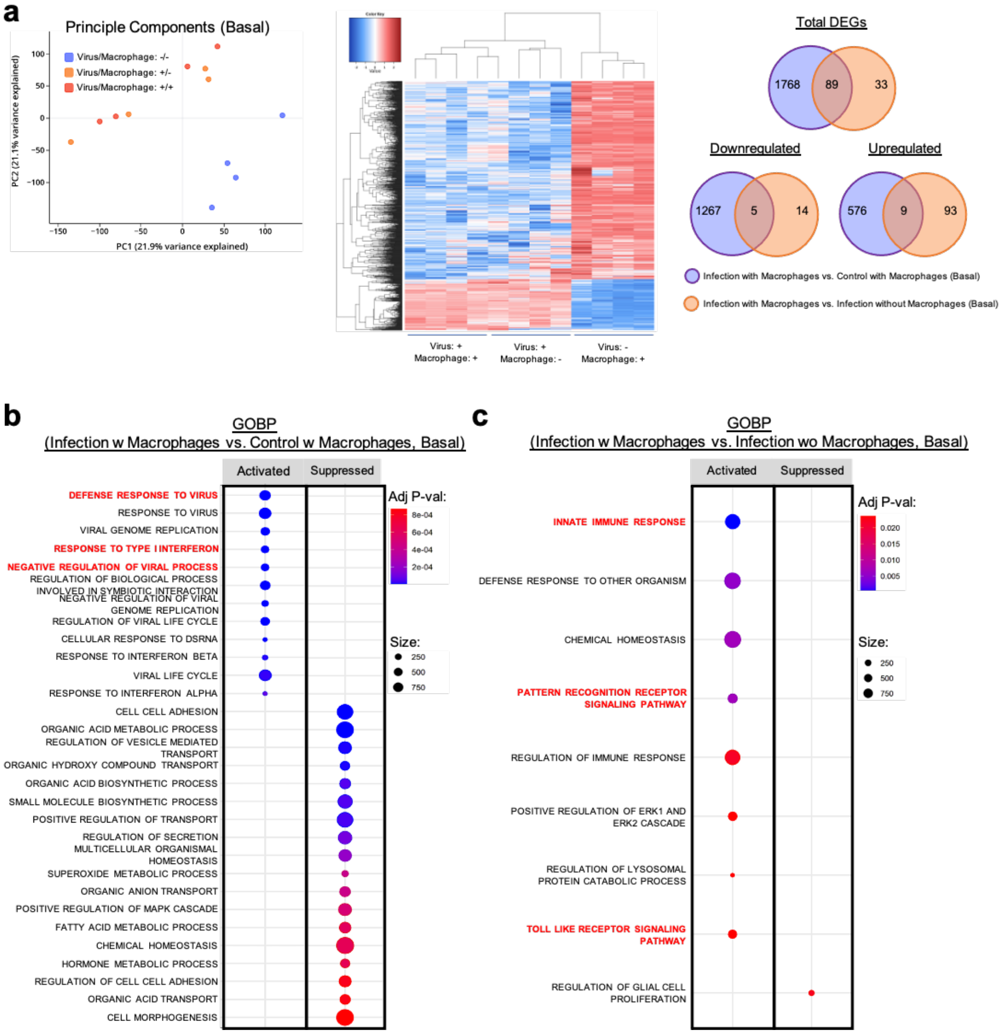


Figure S5. (a) Plot (left) showing principal component analysis of the endothelium with different clusters formed by the three indicated groups. Heatmap (middle) showing clustering analysis of DEGs among the three groups. Venn diagrams (right) depicting the shared or unique DEGs between each comparison. (b) Enriched gene sets revealed by GOBP analysis of DEGs in the endothelium in infected chips compared to mock infected chips, both with macrophages. 30 out of 80 are shown. (c) Enriched gene sets revealed by GOBP analysis of DEGs in the endothelium in infected chips with macrophages compared to infected chips without macrophages. The color of the dots represents the significance of enrichment and the size of the dots represents the input number of each term in (b) and (c).

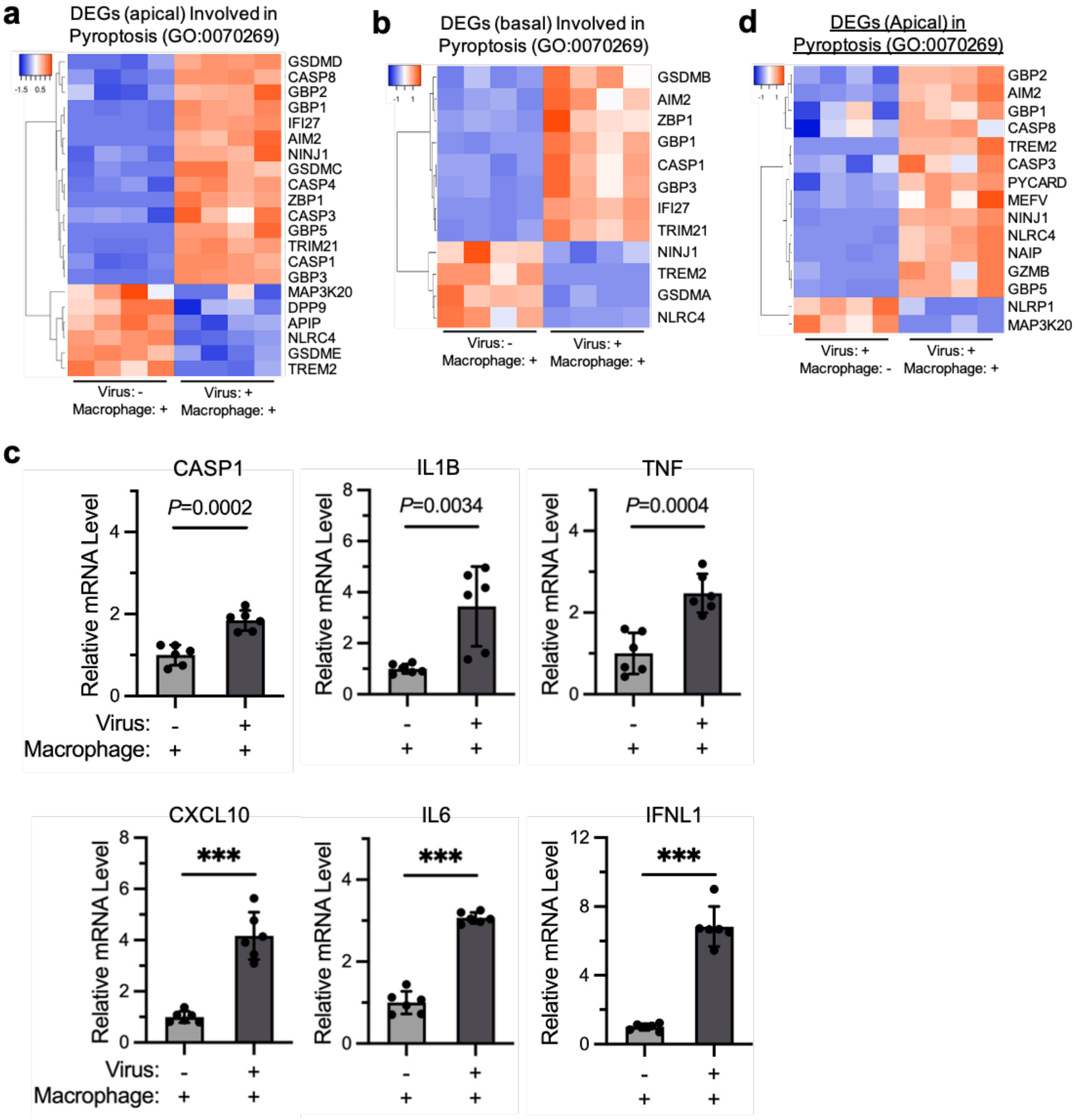


Figure S6. Heatmaps showing DEGs involved in Pyroptosis (GO: 0070269) in samples collected from (a) the apical channels and (b) the basal channels comparing infected chips to mock infected chips, both with macrophages. (c) RT-qPCR analysis of relative mRNA levels of CASP1, IL1B, TNF, CXCL10, IL6 and IFNL1 in the total RNA extracted from the apical channels of infected chips and mock infected chips, both with macrophages. (d) Heatmaps showing DEGs involved in Pyroptosis (GO: 0070269) in samples collected from the apical channels comparing infected chips with macrophages to those without. Data shown are mean ± SD, n = 6 chips in each group in (c); unpaired t test; *** denotes $P < 0.0001$.

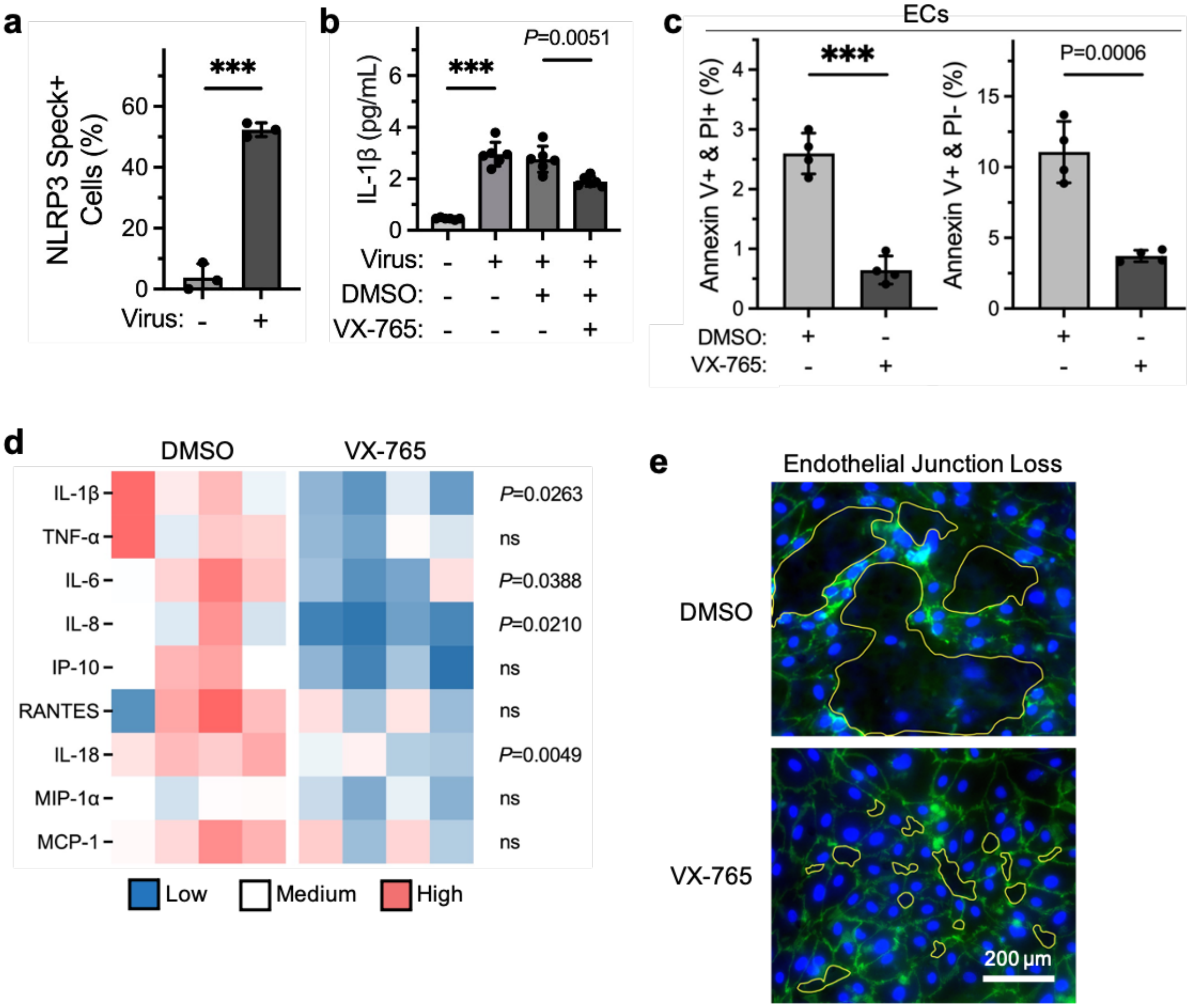


Figure S7. (a) Image quantification of NLRP3 speck$^{+}$ macrophages in submerged culture conditions at 48 h post IAV infection. (b) Supernatant IL-1β levels produced by macrophages in submerged culture conditions at 48 h post IAV infection. (c) Graphs showing that VX-765 treatment significantly reduced both early apoptotic (Annexin V$^{+}$/PI$^{-}$) and late apoptotic (Annexin V$^{+}$/PI$^{+}$) ECs. (d) Heatmap showing apical wash cytokine profiles of chips treated with VX-765 or DMSO at 48 h after IAV infection. (e) Immunofluorescence with VE-cadherin and DAPI showing EC junction loss, as indicated by yellow-outlined area, in chips treated with VX-765 or DMSO at 48 h after IAV infection. Data shown are mean ± SD; n = 3 wells averaged from 3 FOVs per well in each group in (a); n = 6 wells in each group in (b); n = 4 chips in each group in (c) and (d); unpaired t-test; *** denotes $P < 0.0001$.

**Supplementary Tables**

Table S1. GSEA of KEGG pathways of the alveolar tissue in infected chips compared to mock infected chips, both with macrophages (adj $P < 0.001$).

| **Gene set** | **Adjusted *P*-value** | **Normalized enrichment score (NES)** | **Size** |
|---|---|---|---|
| KEGG CYTOKINE CYTOKINE RECEPTOR INTERACTION | 2.112509E-13 | 2.236879 | 214 |
| KEGG RIBOSOME | 1.169284E-9 | -2.44631124166931 | 85 |
| KEGG JAK STAT SIGNALING PATHWAY | 0.000002 | 2.03187387135989 | 125 |
| KEGG CYTOSOLIC DNA SENSING PATHWAY | 0.000008 | 2.1322191220296 | 45 |
| KEGG CHEMOKINE SIGNALING PATHWAY | 0.000175 | 1.79320275889963 | 171 |
| KEGG OXIDATIVE PHOSPHORYLATION | 0.000175 | -1.88893318476036 | 121 |
| KEGG PROPANOATE METABOLISM | 0.000325 | -2.11705498540224 | 30 |
| KEGG SYSTEMIC LUPUS ERYTHEMATOSUS | 0.000577 | 1.86782928353368 | 90 |

Table S2. GSEA of HALLMARK pathways of the alveolar tissue in infected chips compared to mock infected chips, both with macrophages (adj $P < 0.001$).

| **Gene set** | **Adjusted *P*-value** | **NES** | **Size** |
|---|---|---|---|
| HALLMARK INTERFERON GAMMA RESPONSE | 6.562881E-37 | 2.749233 | 196 |
| HALLMARK INTERFERON ALPHA RESPONSE | 1.008182E-25 | 2.713401 | 97 |
| HALLMARK INFLAMMATORY RESPONSE | 6.416873E-10 | 2.069095 | 191 |
| HALLMARK TNFA SIGNALING VIA NFKB | 9.111568E-06 | 1.824120 | 200 |
| HALLMARK OXIDATIVE PHOSPHORYLATION | 0.000047 | -1.74805 | 200 |
| HALLMARK ALLOGRAFT REJECTION | 0.000058 | 1.757308 | 180 |
| HALLMARK IL6 JAK STAT3 SIGNALING | 0.000058 | 1.911650 | 84 |
| HALLMARK FATTY ACID METABOLISM | 0.000077 | -1.75600 | 148 |

Table S3. GOBP analysis of the alveolar tissue in infected chips compared to mock infected chips, both with macrophages (adj $P < 0.001$).

| **Gene set** | **Adjusted *P*-value** | **NES** | **Size** |
|---|---|---|---|
| GOBP DEFENSE RESPONSE TO OTHER ORGANISM | 4.71E-30 | 2.07473534 | 931 |
| GOBP INNATE IMMUNE RESPONSE | 3.07E-27 | 2.10404742 | 766 |
| GOBP CYTOKINE MEDIATED SIGNALING PATHWAY | 1.14E-25 | 2.12529986 | 699 |
| GOBP DEFENSE RESPONSE TO VIRUS | 2.27E-24 | 2.51302567 | 230 |
| GOBP RESPONSE TO VIRUS | 2.65E-21 | 2.36998737 | 317 |
| GOBP RESPONSE TO TYPE I INTERFERON | 1.79E-18 | 2.60271781 | 89 |
| GOBP REGULATION OF IMMUNE RESPONSE | 3.75E-16 | 1.8752238 | 777 |
| GOBP VIRAL GENOME REPLICATION | 5.82E-13 | 2.39549738 | 122 |
| GOBP NEGATIVE REGULATION OF VIRAL PROCESS | 6.24E-13 | 2.45014346 | 81 |
| GOBP REGULATION OF RESPONSE TO BIOTIC STIMULUS | 6.61E-13 | 2.04126899 | 379 |
| GOBP REGULATION OF VIRAL LIFE CYCLE | 9.91E-13 | 2.34529376 | 137 |
| GOBP POSITIVE REGULATION OF IMMUNE SYSTEM PROCESS | 2.20E-12 | 1.76252765 | 843 |
| GOBP POSITIVE REGULATION OF IMMUNE RESPONSE | 4.46E-12 | 1.88473887 | 572 |
| GOBP POSITIVE REGULATION OF RESPONSE TO BIOTIC STIMULUS | 5.65E-12 | 2.14823019 | 233 |
| GOBP RESPONSE TO INTERFERON GAMMA | 5.65E-12 | 2.25022592 | 179 |
| GOBP REGULATION OF BIOLOGICAL PROCESS INVOLVED IN SYMBIOTIC INTERACTION | 9.51E-12 | 2.23079362 | 185 |

| | | | |
|---|---|---|---|
| GOBP NEGATIVE REGULATION OF VIRAL GENOME REPLICATION | 2.90E-11 | 2.45108399 | 52 |
| GOBP REGULATION OF VIRAL GENOME REPLICATION | 2.90E-11 | 2.42657657 | 80 |
| GOBP REGULATION OF RESPONSE TO EXTERNAL STIMULUS | 3.51E-11 | 1.7034944 | 893 |
| GOBP REGULATION OF DEFENSE RESPONSE | 3.41E-10 | 1.79562049 | 590 |
| GOBP CYTOKINE PRODUCTION | 2.58E-09 | 1.75401456 | 665 |
| GOBP REGULATION OF INNATE IMMUNE RESPONSE | 2.78E-09 | 2.01968834 | 285 |
| GOBP RESPONSE TO BACTERIUM | 2.78E-09 | 1.81539897 | 499 |
| GOBP ADAPTIVE IMMUNE RESPONSE | 2.96E-09 | 1.93806098 | 360 |
| GOBP REGULATION OF IMMUNE EFFECTOR PROCESS | 1.09E-08 | 1.91672265 | 354 |
| GOBP KERATINIZATION | 2.44E-08 | 2.22518864 | 98 |
| GOBP KERATINOCYTE DIFFERENTIATION | 4.89E-08 | 2.08022938 | 170 |
| GOBP POSITIVE REGULATION OF DEFENSE RESPONSE | 1.29E-07 | 1.86281879 | 321 |
| GOBP POSITIVE REGULATION OF RESPONSE TO EXTERNAL STIMULUS | 2.66E-07 | 1.786186 | 450 |
| GOBP CORNIFICATION | 3.12E-07 | 2.18190758 | 84 |
| GOBP NEGATIVE REGULATION OF IMMUNE SYSTEM PROCESS | 3.78E-07 | 1.85550926 | 355 |
| GOBP SKIN DEVELOPMENT | 3.78E-07 | 1.91228176 | 272 |
| GOBP INFLAMMATORY RESPONSE | 4.66E-07 | 1.66248943 | 632 |
| GOBP EPIDERMAL CELL DIFFERENTIATION | 4.79E-07 | 1.94454617 | 218 |
| GOBP HUMORAL IMMUNE RESPONSE | 6.15E-07 | 1.9651163 | 192 |
| GOBP ORGANIC ACID METABOLIC PROCESS | 1.07E-06 | -1.4503415 | 930 |
| GOBP VIRAL LIFE CYCLE | 1.31E-06 | 1.81550418 | 329 |

| | | | |
|---|---|---|---|
| GOBP NEGATIVE REGULATION OF LYMPHOCYTE ACTIVATION | 1.43E-06 | 2.03882086 | 142 |
| GOBP RECEPTOR SIGNALING PATHWAY VIA STAT | 2.75E-06 | 2.06330569 | 129 |
| GOBP NEGATIVE REGULATION OF LEUKOCYTE CELL CELL ADHESION | 3.79E-06 | 2.01565823 | 121 |
| GOBP REGULATION OF T CELL ACTIVATION | 4.63E-06 | 1.81339819 | 301 |
| GOBP ADAPTIVE IMMUNE RESPONSE BASED ON SOMATIC RECOMBINATION OF IMMUNE RECEPTORS BUILT FROM IMMUNOGLOBULIN SUPERFAMILY DOMAINS | 5.73E-06 | 1.85610633 | 236 |
| GOBP EPIDERMIS DEVELOPMENT | 6.70E-06 | 1.80012081 | 311 |
| GOBP NEGATIVE REGULATION OF CELL CELL ADHESION | 6.70E-06 | 1.93217406 | 169 |
| GOBP RESPONSE TO INTERFERON BETA | 6.70E-06 | 2.13173261 | 29 |
| GOBP REGULATION OF CELL ACTIVATION | 9.64E-06 | 1.66548219 | 511 |
| GOBP REGULATION OF LYMPHOCYTE ACTIVATION | 9.82E-06 | 1.71157205 | 401 |
| GOBP LYMPHOCYTE MEDIATED IMMUNITY | 1.02E-05 | 1.86274098 | 220 |
| GOBP INTERFERON GAMMA MEDIATED SIGNALING PATHWAY | 1.25E-05 | 2.07829355 | 86 |
| GOBP NEGATIVE REGULATION OF IMMUNE RESPONSE | 1.31E-05 | 1.99292594 | 131 |
| GOBP CELL CHEMOTAXIS | 1.32E-05 | 1.82024258 | 267 |
| GOBP CYTOKINE PRODUCTION INVOLVED IN IMMUNE RESPONSE | 1.71E-05 | 2.07122749 | 86 |
| GOBP RESPONSE TO CHEMOKINE | 1.84E-05 | 2.07506851 | 82 |

| | | | |
|---|---|---|---|
| GOBP NEGATIVE REGULATION OF CELL ACTIVATION | 2.16E-05 | 1.89038082 | 181 |
| GOBP NEGATIVE REGULATION OF MULTICELLULAR ORGANISMAL PROCESS | 2.28E-05 | 1.49607575 | 864 |
| GOBP DEFENSE RESPONSE TO BACTERIUM | 2.32E-05 | 1.89507268 | 178 |
| GOBP MYELOID LEUKOCYTE MIGRATION | 2.48E-05 | 1.84985802 | 194 |
| GOBP ANTIMICROBIAL HUMORAL RESPONSE | 2.78E-05 | 2.02937492 | 92 |
| GOBP COTRANSLATIONAL PROTEIN TARGETING TO MEMBRANE | 2.78E-05 | -2.1010672 | 104 |
| GOBP LEUKOCYTE CELL CELL ADHESION | 2.78E-05 | 1.74532835 | 327 |
| GOBP LYMPHOCYTE ACTIVATION | 2.85E-05 | 1.56884464 | 610 |
| GOBP PROTEIN LOCALIZATION TO ENDOPLASMIC RETICULUM | 2.85E-05 | -1.9733686 | 145 |
| GOBP T CELL ACTIVATION | 2.98E-05 | 1.68380826 | 425 |
| GOBP NEGATIVE REGULATION OF CYTOKINE PRODUCTION | 3.58E-05 | 1.78562164 | 258 |
| GOBP IMMUNE SYSTEM DEVELOPMENT | 3.81E-05 | 1.48913879 | 885 |
| GOBP REGULATION OF LEUKOCYTE MEDIATED IMMUNITY | 3.95E-05 | 1.85026791 | 190 |
| GOBP ESTABLISHMENT OF PROTEIN LOCALIZATION TO ENDOPLASMIC RETICULUM | 5.14E-05 | -1.995198 | 118 |
| GOBP POSITIVE REGULATION OF CYTOKINE PRODUCTION | 6.73E-05 | 1.66880999 | 386 |
| GOBP LEUKOCYTE CHEMOTAXIS | 9.43E-05 | 1.78656122 | 200 |
| GOBP NEGATIVE REGULATION OF CELL ADHESION | 9.72E-05 | 1.7775897 | 262 |

| | | | |
|---|---|---|---|
| GOBP CELLULAR LIPID METABOLIC PROCESS | 9.92E-05 | -1.373471 | 924 |
| GOBP CELLULAR RESPONSE TO MOLECULE OF BACTERIAL ORIGIN | 9.98E-05 | 1.85299534 | 179 |
| GOBP REGULATION OF ADAPTIVE IMMUNE RESPONSE | 0.00011897 | 1.84424276 | 160 |
| GOBP CELLULAR RESPONSE TO BIOTIC STIMULUS | 0.00012107 | 1.77643382 | 203 |
| GOBP REGULATION OF LYMPHOCYTE MEDIATED IMMUNITY | 0.00012107 | 1.90384148 | 140 |
| GOBP POSITIVE REGULATION OF LEUKOCYTE MEDIATED IMMUNITY | 0.00012651 | 1.95011942 | 112 |
| GOBP REGULATION OF CELL CELL ADHESION | 0.00016111 | 1.64425703 | 397 |
| GOBP BLOOD VESSEL MORPHOGENESIS | 0.00020183 | 1.54044064 | 548 |
| GOBP MICROTUBULE BUNDLE FORMATION | 0.00023441 | -1.9746998 | 99 |
| GOBP RESPONSE TO MOLECULE OF BACTERIAL ORIGIN | 0.00024974 | 1.67567914 | 300 |
| GOBP T CELL DIFFERENTIATION | 0.00028225 | 1.7264325 | 227 |
| GOBP CELL KILLING | 0.0002875 | 1.86246226 | 134 |
| GOBP NEUTROPHIL MIGRATION | 0.00032265 | 1.90642193 | 109 |
| GOBP GRANULOCYTE MIGRATION | 0.00033085 | 1.86595975 | 131 |
| GOBP CILIUM ORGANIZATION | 0.00034608 | -1.5781682 | 387 |
| GOBP REGULATION OF PRODUCTION OF MOLECULAR MEDIATOR OF IMMUNE RESPONSE | 0.00046494 | 1.88458405 | 130 |
| GOBP REGULATION OF DEFENSE RESPONSE TO VIRUS | 0.00047421 | 1.94894787 | 65 |
| GOBP LYMPHOCYTE MIGRATION | 0.00050765 | 1.90076661 | 101 |
| GOBP ANTIMICROBIAL HUMORAL IMMUNE RESPONSE MEDIATED | 0.0005091 | 2.00226841 | 51 |

| | | | |
|---|---|---|---|
| BY ANTIMICROBIAL PEPTIDE | | | |
| GOBP FATTY ACID DERIVATIVE BIOSYNTHETIC PROCESS | 0.0005091 | -2.0773045 | 57 |
| GOBP REGULATION OF CELL ADHESION | 0.0005091 | 1.48518886 | 669 |
| GOBP REGULATION OF RESPONSE TO CYTOKINE STIMULUS | 0.0005091 | 1.81536388 | 163 |
| GOBP TAXIS | 0.0005091 | 1.53235918 | 569 |
| GOBP T CELL MEDIATED IMMUNITY | 0.00051297 | 1.91718691 | 92 |
| GOBP TRANSLATIONAL INITIATION | 0.00052372 | -1.7586546 | 185 |
| GOBP LEUKOTRIENE BIOSYNTHETIC PROCESS | 0.00053461 | -2.0818937 | 18 |
| GOBP SMALL MOLECULE BIOSYNTHETIC PROCESS | 0.00054726 | -1.4486475 | 606 |
| GOBP MONONUCLEAR CELL DIFFERENTIATION | 0.00056401 | 1.60467849 | 369 |
| GOBP G PROTEIN COUPLED RECEPTOR SIGNALING PATHWAY | 0.00058366 | 1.48390528 | 667 |
| GOBP LYMPHOCYTE CHEMOTAXIS | 0.00061698 | 2.03172116 | 50 |
| GOBP ANIMAL ORGAN MORPHOGENESIS | 0.0006287 | 1.42449246 | 872 |
| GOBP POSITIVE REGULATION OF NATURAL KILLER CELL MEDIATED IMMUNITY | 0.0006287 | 1.96505539 | 25 |
| GOBP MONOCARBOXYLIC ACID METABOLIC PROCESS | 0.00065092 | -1.4513354 | 547 |
| GOBP MONONUCLEAR CELL MIGRATION | 0.00065092 | 1.76728313 | 167 |
| GOBP RESPONSE TO DSRNA | 0.00065092 | 2.01860329 | 42 |
| GOBP LEUKOCYTE MIGRATION | 0.00066395 | 1.59080673 | 398 |
| GOBP CELLULAR RESPIRATION | 0.00067027 | -1.7173162 | 175 |
| GOBP POSITIVE REGULATION OF IMMUNE EFFECTOR PROCESS | 0.00067027 | 1.76213684 | 191 |
| GOBP FATTY ACID METABOLIC PROCESS | 0.00069321 | -1.5796259 | 336 |

| | | | |
|---|---|---|---|
| GOBP TUBE MORPHOGENESIS | 0.00071418 | 1.44091969 | 750 |
| GOBP VASCULATURE DEVELOPMENT | 0.00078229 | 1.46166584 | 644 |
| GOBP REGULATION OF RIBONUCLEASE ACTIVITY | 0.0008528 | 1.88848115 | 9 |
| GOBP NATURAL KILLER CELL MEDIATED IMMUNITY | 0.00086121 | 1.98809194 | 55 |
| GOBP ORGANIC ACID BIOSYNTHETIC PROCESS | 0.00089076 | -1.6027658 | 278 |
| GOBP REGULATION OF NATURAL KILLER CELL MEDIATED IMMUNITY | 0.00090072 | 1.97540403 | 39 |
| GOBP OXIDATIVE PHOSPHORYLATION | 0.00096947 | -1.7971707 | 136 |
| GOBP MONOCYTE CHEMOTAXIS | 0.00098961 | 1.97922857 | 55 |

Table S4. GSEA of KEGG pathways of the alveolar tissue in infected chips with macrophages compared to infected chips without macrophages (adj $P < 0.001$).

| **Gene set** | **Adjusted *P*-value** | **NES** | **Size** |
|---|---|---|---|
| KEGG RIBOSOME | 9.954384E-13 | -2.613737 | 86 |
| KEGG AUTOIMMUNE THYROID DISEASE | 5.811410E-9 | 2.2131192 | 32 |
| KEGG CELL CYCLE | 5.811410E-9 | -2.269658 | 122 |
| KEGG SYSTEMIC LUPUS ERYTHEMATOSUS | 5.811410E-9 | 2.150499 | 94 |
| KEGG LYSOSOME | 5.811410E-9 | 2.092364 | 120 |
| KEGG CHEMOKINE SIGNALING PATHWAY | 4.236431E-8 | 1.977527 | 173 |
| KEGG LEISHMANIA INFECTION | 4.236431E-8 | 2.180616 | 67 |
| KEGG ANTIGEN PROCESSING AND PRESENTATION | 7.283604E-8 | 2.168275 | 60 |
| KEGG CYTOKINE CYTOKINE RECEPTOR INTERACTION | 1.393202E-7 | 1.877050 | 213 |
| KEGG NATURAL KILLER CELL MEDIATED CYTOTOXICITY | 2.573652E-7 | 2.024443 | 105 |
| KEGG FC GAMMA R MEDIATED PHAGOCYTOSIS | 0.000001 | 2.010858 | 96 |
| KEGG DNA REPLICATION | 0.000002 | -2.32282 | 36 |
| KEGG ALLOGRAFT REJECTION | 0.000003 | 2.055974 | 28 |
| KEGG B CELL RECEPTOR SIGNALING PATHWAY | 0.000003 | 2.005037 | 73 |
| KEGG TOLL LIKE RECEPTOR SIGNALING PATHWAY | 0.000006 | 1.996577 | 91 |
| KEGG HEMATOPOIETIC CELL LINEAGE | 0.000009 | 1.979756 | 75 |
| KEGG SPLICEOSOME | 0.000024 | -1.89824 | 126 |
| KEGG VIRAL MYOCARDITIS | 0.000073 | 1.974961 | 60 |
| KEGG CELL ADHESION MOLECULES CAMS | 0.000084 | 1.819725 | 122 |
| KEGG TYPE I DIABETES MELLITUS | 0.000093 | 1.941452 | 35 |
| KEGG INTESTINAL IMMUNE NETWORK FOR IGA PRODUCTION | 0.000108 | 1.948771 | 34 |
| KEGG OOCYTE MEIOSIS | 0.000293 | -1.79681 | 102 |

| | | | |
|---|---|---|---|
| KEGG GRAFT VERSUS HOST DISEASE | 0.000387 | 1.925607 | 29 |
| KEGG LEUKOCYTE TRANSENDOTHELIAL MIGRATION | 0.000540 | 1.733327 | 107 |
| KEGG COMPLEMENT AND COAGULATION CASCADES | 0.000705 | 1.835839 | 59 |
| KEGG ASTHMA | 0.000948 | 1.845972 | 19 |

Table S5. GSEA of HALLMARK pathways of the alveolar tissue in infected chips with macrophages compared to infected chips without macrophages (adj $P < 0.001$).

| **Gene set** | **Adjusted *P*-value** | **NES** | **Size** |
|---|---|---|---|
| HALLMARK E2F TARGETS | 2.613499E-32 | -2.92036 | 200 |
| HALLMARK G2M CHECKPOINT | 1.529239E-28 | -0.67201 | 198 |
| HALLMARK MYC TARGETS V1 | 1.745885E-23 | -2.66867 | 199 |
| HALLMARK EPITHELIAL MESENCHYMAL TRANSITION | 8.629709E-17 | -2.42495 | 199 |
| HALLMARK ALLOGRAFT REJECTION | 1.357807E-15 | 2.235692 | 183 |
| HALLMARK MYC TARGETS V2 | 1.600264E-10 | -2.52652 | 57 |
| HALLMARK INTERFERON GAMMA RESPONSE | 5.164594E-10 | 2.01768 | 196 |
| HALLMARK INFLAMMATORY RESPONSE | 2.144777E-08 | 1.94342 | 189 |
| HALLMARK COMPLEMENT | 5.9670009E-06 | 1.80267 | 191 |
| HALLMARK MITOTIC SPINDLE | 0.0000105 | -1.6915 | 199 |
| HALLMARK INTERFERON ALPHA RESPONSE | 0.0000105 | 1.90170 | 97 |
| HALLMARK IL2 STAT5 SIGNALING | 0.0000680 | 1.71518 | 194 |
| HALLMARK KRAS SIGNALING UP | 0.0001170 | 1.67505 | 192 |

Table S6. GOBP analysis of the alveolar tissue in infected chips with macrophages compared to infected chips without macrophages (adj $P < 0.001$).

| Gene set | Adjusted *P*-value | NES | Size |
|---|---|---|---|
| GOBP REGULATION OF IMMUNE RESPONSE | 3.31E-55 | 2.24277938 | 776 |
| GOBP POSITIVE REGULATION OF IMMUNE SYSTEM PROCESS | 1.67E-46 | 2.13332965 | 841 |
| GOBP DEFENSE RESPONSE TO OTHER ORGANISM | 1.31E-44 | 2.08077975 | 926 |
| GOBP INNATE IMMUNE RESPONSE | 5.40E-43 | 2.13849936 | 765 |
| GOBP LEUKOCYTE MEDIATED IMMUNITY | 1.82E-41 | 2.15887867 | 711 |
| GOBP CELL ACTIVATION INVOLVED IN IMMUNE RESPONSE | 3.34E-40 | 2.16895072 | 664 |
| GOBP MYELOID LEUKOCYTE ACTIVATION | 3.84E-38 | 2.17360269 | 619 |
| GOBP POSITIVE REGULATION OF IMMUNE RESPONSE | 6.03E-38 | 2.20386603 | 573 |
| GOBP LYMPHOCYTE ACTIVATION | 2.12E-32 | 2.10163681 | 606 |
| GOBP REGULATION OF CELL ACTIVATION | 1.22E-31 | 2.16445554 | 509 |
| GOBP ADAPTIVE IMMUNE RESPONSE | 4.39E-29 | 2.2780062 | 360 |
| GOBP ACTIVATION OF IMMUNE RESPONSE | 1.56E-28 | 2.21374598 | 409 |
| GOBP MYELOID LEUKOCYTE MEDIATED IMMUNITY | 5.30E-28 | 2.10947677 | 518 |
| GOBP IMMUNE RESPONSE REGULATING SIGNALING PATHWAY | 1.15E-26 | 2.20816019 | 369 |
| GOBP EXOCYTOSIS | 3.47E-26 | 1.91635047 | 838 |
| GOBP REGULATION OF IMMUNE EFFECTOR PROCESS | 1.34E-25 | 2.21393677 | 353 |
| GOBP T CELL ACTIVATION | 2.63E-25 | 2.13921075 | 423 |
| GOBP REGULATION OF LYMPHOCYTE ACTIVATION | 1.13E-23 | 2.12630481 | 397 |

| | | | |
|---|---|---|---|
| GOBP DNA REPLICATION | 1.44E-23 | -2.7112216 | 271 |
| GOBP INFLAMMATORY RESPONSE | 4.36E-23 | 1.96552253 | 637 |
| GOBP REGULATION OF RESPONSE TO EXTERNAL STIMULUS | 1.35E-22 | 1.83379899 | 889 |
| GOBP LEUKOCYTE PROLIFERATION | 1.43E-21 | 2.23156038 | 272 |
| GOBP CYTOKINE PRODUCTION | 1.55E-21 | 1.90775821 | 666 |
| GOBP RIBOSOME BIOGENESIS | 4.05E-21 | -2.5639436 | 299 |
| GOBP LEUKOCYTE DIFFERENTIATION | 5.10E-21 | 2.02414474 | 468 |
| GOBP CHROMOSOME SEGREGATION | 7.46E-21 | -2.5148551 | 314 |
| GOBP REGULATION OF T CELL ACTIVATION | 2.00E-19 | 2.15288209 | 299 |
| GOBP NCRNA PROCESSING | 4.65E-19 | -2.3150561 | 381 |
| GOBP POSITIVE REGULATION OF CELL ACTIVATION | 4.89E-19 | 2.14740654 | 304 |
| GOBP MITOTIC NUCLEAR DIVISION | 7.15E-19 | -2.5109345 | 284 |
| GOBP PHAGOCYTOSIS | 7.15E-19 | 2.20221448 | 250 |
| GOBP REGULATION OF DEFENSE RESPONSE | 2.00E-18 | 1.90412051 | 586 |
| GOBP NEGATIVE REGULATION OF IMMUNE SYSTEM PROCESS | 3.25E-18 | 2.05395048 | 354 |
| GOBP LEUKOCYTE CELL CELL ADHESION | 4.15E-18 | 2.09898119 | 324 |
| GOBP NUCLEAR CHROMOSOME SEGREGATION | 1.17E-17 | -2.5026278 | 253 |
| GOBP POSITIVE REGULATION OF CYTOKINE PRODUCTION | 1.61E-17 | 2.03917273 | 384 |
| GOBP DNA DEPENDENT DNA REPLICATION | 3.01E-17 | -2.7017278 | 153 |
| GOBP REGULATION OF CELL CELL ADHESION | 3.01E-17 | 2.00439341 | 393 |
| GOBP REGULATION OF LEUKOCYTE PROLIFERATION | 3.18E-17 | 2.23556749 | 215 |
| GOBP MONONUCLEAR CELL DIFFERENTIATION | 8.97E-17 | 2.02144275 | 367 |

| | | | |
|---|---|---|---|
| GOBP LEUKOCYTE MIGRATION | 1.47E-16 | 1.97552627 | 397 |
| GOBP MITOTIC SISTER CHROMATID SEGREGATION | 2.24E-16 | -2.6719814 | 160 |
| GOBP POSITIVE REGULATION OF RESPONSE TO EXTERNAL STIMULUS | 3.90E-16 | 1.93685768 | 446 |
| GOBP REGULATION OF LEUKOCYTE MEDIATED IMMUNITY | 4.83E-16 | 2.21541776 | 190 |
| GOBP SISTER CHROMATID SEGREGATION | 1.32E-15 | -2.5827607 | 192 |
| GOBP ADAPTIVE IMMUNE RESPONSE BASED ON SOMATIC RECOMBINATION OF IMMUNE RECEPTORS BUILT FROM IMMUNOGLOBULIN SUPERFAMILY DOMAINS | 3.48E-15 | 2.13015327 | 235 |
| GOBP REGULATION OF RESPONSE TO BIOTIC STIMULUS | 5.66E-15 | 1.97105614 | 376 |
| GOBP T CELL PROLIFERATION | 1.44E-14 | 2.20101576 | 180 |
| GOBP REGULATION OF LEUKOCYTE DIFFERENTIATION | 2.06E-14 | 2.08718552 | 250 |
| GOBP RRNA METABOLIC PROCESS | 2.25E-14 | -2.4515039 | 234 |
| GOBP NEGATIVE REGULATION OF LYMPHOCYTE ACTIVATION | 2.82E-14 | 2.26242583 | 143 |
| GOBP CYTOKINE MEDIATED SIGNALING PATHWAY | 4.06E-14 | 1.7432364 | 704 |
| GOBP LYMPHOCYTE MEDIATED IMMUNITY | 4.87E-14 | 2.11851727 | 218 |
| GOBP NEGATIVE REGULATION OF CELL ACTIVATION | 1.21E-13 | 2.16357128 | 183 |
| GOBP REGULATION OF INNATE IMMUNE RESPONSE | 1.58E-13 | 2.04077294 | 282 |

| | | | |
|---|---|---|---|
| GOBP ANTIGEN RECEPTOR MEDIATED SIGNALING PATHWAY | 2.38E-13 | 2.06523555 | 230 |
| GOBP POSITIVE REGULATION OF CELL CELL ADHESION | 2.96E-13 | 2.06363357 | 249 |
| GOBP POSITIVE REGULATION OF IMMUNE EFFECTOR PROCESS | 7.22E-13 | 2.13399317 | 191 |
| GOBP RESPONSE TO INTERFERON GAMMA | 7.72E-13 | 2.15019744 | 180 |
| GOBP PATTERN RECOGNITION RECEPTOR SIGNALING PATHWAY | 8.05E-13 | 2.13542904 | 193 |
| GOBP IMMUNE SYSTEM DEVELOPMENT | 1.62E-12 | 1.6524796 | 885 |
| GOBP MONONUCLEAR CELL MIGRATION | 1.90E-12 | 2.1338467 | 166 |
| GOBP REGULATION OF CELL ADHESION | 2.56E-12 | 1.72134956 | 665 |
| GOBP ANTIGEN PROCESSING AND PRESENTATION | 2.65E-12 | 2.07037579 | 224 |
| GOBP NEGATIVE REGULATION OF LEUKOCYTE CELL CELL ADHESION | 3.20E-12 | 2.2172694 | 122 |
| GOBP POSITIVE REGULATION OF DEFENSE RESPONSE | 4.16E-12 | 1.94292878 | 318 |
| GOBP T CELL DIFFERENTIATION | 5.28E-12 | 2.05932408 | 225 |
| GOBP CELL KILLING | 5.65E-12 | 2.19326037 | 132 |
| GOBP REGULATION OF HEMOPOIESIS | 6.19E-12 | 1.88213984 | 367 |
| GOBP TOLL LIKE RECEPTOR SIGNALING PATHWAY | 6.19E-12 | 2.16182611 | 145 |
| GOBP NEGATIVE REGULATION OF LEUKOCYTE PROLIFERATION | 6.55E-12 | 2.28473265 | 83 |
| GOBP REGULATION OF ADAPTIVE IMMUNE RESPONSE | 9.21E-12 | 2.12201392 | 159 |
| GOBP REGULATION OF LYMPHOCYTE MEDIATED IMMUNITY | 1.09E-11 | 2.16091599 | 138 |

| | | | |
|---|---|---|---|
| GOBP DNA CONFORMATION CHANGE | 1.70E-11 | -2.1107372 | 301 |
| GOBP POSITIVE REGULATION OF LEUKOCYTE CELL CELL ADHESION | 1.75E-11 | 2.06411562 | 209 |
| GOBP RESPONSE TO BACTERIUM | 1.86E-11 | 1.77729282 | 502 |
| GOBP LYMPHOCYTE ACTIVATION INVOLVED IN IMMUNE RESPONSE | 3.33E-11 | 2.12436667 | 161 |
| GOBP POSITIVE REGULATION OF ERK1 AND ERK2 CASCADE | 6.62E-11 | 2.0909446 | 168 |
| GOBP LEUKOCYTE CHEMOTAXIS | 7.12E-11 | 2.04974719 | 200 |
| GOBP POSITIVE REGULATION OF CELL ADHESION | 1.07E-10 | 1.82293911 | 390 |
| GOBP POSITIVE REGULATION OF MAPK CASCADE | 1.08E-10 | 1.77785657 | 458 |
| GOBP MEIOTIC CELL CYCLE | 1.17E-10 | -2.2987623 | 207 |
| GOBP RIBOSOMAL LARGE SUBUNIT BIOGENESIS | 1.20E-10 | -2.5954958 | 70 |
| GOBP T CELL ACTIVATION INVOLVED IN IMMUNE RESPONSE | 1.32E-10 | 2.19342127 | 90 |
| GOBP REGULATION OF VESICLE MEDIATED TRANSPORT | 1.77E-10 | 1.76048477 | 471 |
| GOBP NUCLEAR TRANSCRIBED MRNA CATABOLIC PROCESS NONSENSE MEDIATED DECAY | 1.80E-10 | -2.4451779 | 120 |
| GOBP CELL CELL ADHESION | 2.09E-10 | 1.62997352 | 784 |
| GOBP POSITIVE REGULATION OF RESPONSE TO BIOTIC STIMULUS | 2.13E-10 | 1.96171909 | 230 |
| GOBP TUMOR NECROSIS FACTOR SUPERFAMILY CYTOKINE PRODUCTION | 2.31E-10 | 2.11527903 | 133 |

| | | | |
|---|---|---|---|
| GOBP POSITIVE REGULATION OF HEMOPOIESIS | 2.35E-10 | 2.10709703 | 138 |
| GOBP MITOCHONDRIAL GENE EXPRESSION | 2.42E-10 | -2.319175 | 165 |
| GOBP MITOTIC CELL CYCLE CHECKPOINT | 2.44E-10 | -2.3308184 | 153 |
| GOBP ALPHA BETA T CELL ACTIVATION | 2.71E-10 | 2.11346761 | 133 |
| GOBP MYELOID LEUKOCYTE MIGRATION | 4.68E-10 | 2.01810277 | 196 |
| GOBP NEGATIVE REGULATION OF CELL CELL ADHESION | 5.16E-10 | 2.05857542 | 170 |
| GOBP REGULATION OF LYMPHOCYTE DIFFERENTIATION | 5.18E-10 | 2.0769638 | 161 |
| GOBP REGULATION OF NUCLEAR DIVISION | 6.27E-10 | -2.4028199 | 120 |
| GOBP REGULATION OF MITOTIC NUCLEAR DIVISION | 6.77E-10 | -2.4852112 | 103 |
| GOBP B CELL RECEPTOR SIGNALING PATHWAY | 1.28E-09 | 2.26248358 | 57 |
| GOBP CELL CYCLE CHECKPOINT | 1.28E-09 | -2.2145772 | 202 |
| GOBP SPINDLE ORGANIZATION | 1.28E-09 | -2.2633708 | 179 |
| GOBP CELL CYCLE G2 M PHASE TRANSITION | 1.65E-09 | -2.0749308 | 264 |
| GOBP ESTABLISHMENT OF PROTEIN LOCALIZATION TO ENDOPLASMIC RETICULUM | 1.82E-09 | -2.3623198 | 118 |
| GOBP TRANSLATIONAL INITIATION | 1.85E-09 | -2.2554482 | 186 |
| GOBP HUMORAL IMMUNE RESPONSE | 1.90E-09 | 1.9983098 | 192 |
| GOBP MEIOTIC CELL CYCLE PROCESS | 1.90E-09 | -2.2528549 | 156 |
| GOBP REGULATION OF CELL KILLING | 1.92E-09 | 2.19715142 | 82 |
| GOBP MEMBRANE INVAGINATION | 2.02E-09 | 2.22389809 | 68 |
| GOBP REGULATION OF MAPK CASCADE | 2.02E-09 | 1.64059432 | 617 |
| GOBP ERK1 AND ERK2 CASCADE | 2.14E-09 | 1.92130417 | 265 |

| | | | |
|---|---|---|---|
| GOBP COTRANSLATIONAL PROTEIN TARGETING TO MEMBRANE | 2.46E-09 | -2.4039471 | 104 |
| GOBP NEGATIVE REGULATION OF T CELL PROLIFERATION | 2.79E-09 | 2.21627677 | 64 |
| GOBP POSITIVE REGULATION OF INTRACELLULAR SIGNAL TRANSDUCTION | 3.08E-09 | 1.54922746 | 891 |
| GOBP DOUBLE STRAND BREAK REPAIR | 3.17E-09 | -2.0110896 | 252 |
| GOBP LEUKOCYTE MEDIATED CYTOTOXICITY | 3.55E-09 | 2.14883117 | 92 |
| GOBP NEGATIVE REGULATION OF IMMUNE RESPONSE | 3.55E-09 | 2.06350215 | 130 |
| GOBP MITOCHONDRIAL TRANSLATION | 3.85E-09 | -2.2993635 | 134 |
| GOBP REGULATION OF INFLAMMATORY RESPONSE | 4.27E-09 | 1.86622043 | 289 |
| GOBP B CELL ACTIVATION | 4.56E-09 | 1.94028757 | 222 |
| GOBP MAPK CASCADE | 5.94E-09 | 1.57041817 | 808 |
| GOBP POSITIVE REGULATION OF LEUKOCYTE PROLIFERATION | 6.02E-09 | 2.061944 | 128 |
| GOBP MACROPHAGE ACTIVATION | 6.05E-09 | 2.11151152 | 93 |
| GOBP REGULATION OF T CELL DIFFERENTIATION | 7.21E-09 | 2.05508153 | 134 |
| GOBP MICROTUBULE CYTOSKELETON ORGANIZATION INVOLVED IN MITOSIS | 7.22E-09 | -2.2189103 | 142 |
| GOBP G PROTEIN COUPLED RECEPTOR SIGNALING PATHWAY | 7.34E-09 | 1.60605124 | 686 |
| GOBP INTERFERON GAMMA PRODUCTION | 8.63E-09 | 2.09293274 | 100 |
| GOBP CELL CHEMOTAXIS | 8.91E-09 | 1.88326906 | 268 |
| GOBP REGULATION OF HUMORAL IMMUNE RESPONSE | 1.25E-08 | 2.20054678 | 60 |

| | | | |
|---|---|---|---|
| GOBP LYMPHOCYTE MIGRATION | 1.52E-08 | 2.09770046 | 101 |
| GOBP RIBONUCLEOPROTEIN COMPLEX SUBUNIT ORGANIZATION | 1.90E-08 | -2.2017126 | 194 |
| GOBP POSITIVE REGULATION OF CELL CYCLE PROCESS | 2.04E-08 | -1.9790346 | 257 |
| GOBP POSITIVE REGULATION OF HYDROLASE ACTIVITY | 2.04E-08 | 1.58101351 | 715 |
| GOBP CHROMOSOME SEPARATION | 2.31E-08 | -2.3626242 | 89 |
| GOBP NEGATIVE REGULATION OF CELL ADHESION | 2.32E-08 | 1.86201267 | 263 |
| GOBP INTERLEUKIN 1 PRODUCTION | 2.45E-08 | 2.09133024 | 96 |
| GOBP REGULATION OF CHROMOSOME ORGANIZATION | 2.45E-08 | -1.9945629 | 264 |
| GOBP NEGATIVE REGULATION OF CYTOKINE PRODUCTION | 2.51E-08 | 1.86301954 | 258 |
| GOBP REGULATION OF B CELL ACTIVATION | 2.51E-08 | 2.07651941 | 111 |
| GOBP NEGATIVE REGULATION OF CHROMOSOME ORGANIZATION | 2.54E-08 | -2.3664129 | 87 |
| GOBP POSITIVE REGULATION OF LEUKOCYTE MEDIATED IMMUNITY | 2.82E-08 | 2.07359088 | 111 |
| GOBP GRANULOCYTE MIGRATION | 2.91E-08 | 2.02291476 | 133 |
| GOBP INTERLEUKIN 12 PRODUCTION | 2.91E-08 | 2.19293468 | 53 |
| GOBP REGULATION OF PHAGOCYTOSIS | 3.12E-08 | 2.12478632 | 81 |
| GOBP REGULATION OF CHROMOSOME SEPARATION | 4.41E-08 | -2.4605795 | 67 |
| GOBP B CELL PROLIFERATION | 4.67E-08 | 2.13547526 | 74 |
| GOBP TAXIS | 4.96E-08 | 1.63300263 | 569 |
| GOBP RIBOSOMAL SMALL SUBUNIT BIOGENESIS | 5.54E-08 | -2.4261369 | 72 |

| | | | |
|---|---|---|---|
| GOBP DEFENSE RESPONSE TO BACTERIUM | 5.65E-08 | 1.95404854 | 177 |
| GOBP POSITIVE REGULATION OF LYMPHOCYTE DIFFERENTIATION | 6.62E-08 | 2.08982647 | 92 |
| GOBP DIVALENT INORGANIC CATION HOMEOSTASIS | 7.67E-08 | 1.70283943 | 398 |
| GOBP REGULATION OF CHROMOSOME SEGREGATION | 7.84E-08 | -2.3394629 | 82 |
| GOBP REGULATION OF LEUKOCYTE MEDIATED CYTOTOXICITY | 9.47E-08 | 2.14865099 | 66 |
| GOBP NEGATIVE REGULATION OF CELL CYCLE PROCESS | 9.83E-08 | -1.8478298 | 317 |
| GOBP METAPHASE ANAPHASE TRANSITION OF CELL CYCLE | 1.04E-07 | -2.4238276 | 62 |
| GOBP POSITIVE REGULATION OF T CELL PROLIFERATION | 1.05E-07 | 2.0559576 | 87 |
| GOBP CELL CYCLE DNA REPLICATION | 1.05E-07 | -2.4023739 | 63 |
| GOBP MESENCHYME DEVELOPMENT | 1.10E-07 | -1.9774008 | 247 |
| GOBP PROTEIN LOCALIZATION TO ENDOPLASMIC RETICULUM | 1.10E-07 | -2.0997688 | 145 |
| GOBP METAL ION HOMEOSTASIS | 1.18E-07 | 1.62581074 | 524 |
| GOBP REGULATION OF CELL CYCLE G2 M PHASE TRANSITION | 1.27E-07 | -2.0656678 | 210 |
| GOBP DNA GEOMETRIC CHANGE | 1.34E-07 | -2.233091 | 113 |
| GOBP REGULATION OF ANTIGEN RECEPTOR MEDIATED SIGNALING PATHWAY | 1.41E-07 | 2.15173457 | 60 |
| GOBP REGULATION OF MITOTIC SISTER CHROMATID SEGREGATION | 1.41E-07 | -2.515869 | 43 |
| GOBP NEGATIVE REGULATION OF CELL | 1.53E-07 | -1.9547063 | 236 |

| | | | |
|---|---|---|---|
| CYCLE PHASE TRANSITION | | | |
| GOBP NUCLEAR TRANSCRIBED MRNA CATABOLIC PROCESS | 1.56E-07 | -2.0780041 | 206 |
| GOBP CELLULAR ION HOMEOSTASIS | 1.90E-07 | 1.62324129 | 548 |
| GOBP ION HOMEOSTASIS | 2.17E-07 | 1.56384144 | 649 |
| GOBP POSITIVE REGULATION OF INTERFERON GAMMA PRODUCTION | 2.38E-07 | 2.13726172 | 58 |
| GOBP TRANSLATIONAL TERMINATION | 2.38E-07 | -2.2357862 | 104 |
| GOBP FC RECEPTOR MEDIATED STIMULATORY SIGNALING PATHWAY | 2.48E-07 | 2.06931974 | 83 |
| GOBP CELL CYCLE G1 S PHASE TRANSITION | 2.53E-07 | -1.8923496 | 251 |
| GOBP ALPHA BETA T CELL DIFFERENTIATION | 2.69E-07 | 2.04014455 | 95 |
| GOBP POSITIVE REGULATION OF CELL KILLING | 2.88E-07 | 2.1288626 | 51 |
| GOBP MITOCHONDRIAL TRANSLATIONAL TERMINATION | 3.31E-07 | -2.2613082 | 89 |
| GOBP EMBRYONIC ORGAN DEVELOPMENT | 3.39E-07 | -1.6989679 | 368 |
| GOBP RESPIRATORY BURST | 3.41E-07 | 2.152226 | 32 |
| GOBP NEGATIVE REGULATION OF METAPHASE ANAPHASE TRANSITION OF CELL CYCLE | 3.47E-07 | -2.4543723 | 40 |
| GOBP METAL ION TRANSPORT | 3.83E-07 | 1.66906884 | 400 |
| GOBP POSITIVE REGULATION OF CELL CYCLE | 4.08E-07 | -1.7728195 | 344 |
| GOBP ENDOCYTOSIS | 4.38E-07 | 1.60916398 | 519 |
| GOBP RECOMBINATIONAL REPAIR | 5.13E-07 | -2.0919134 | 136 |
| GOBP NEGATIVE REGULATION OF | 5.79E-07 | 1.98342519 | 113 |

| | | | |
|---|---|---|---|
| IMMUNE EFFECTOR PROCESS | | | |
| GOBP MITOTIC SPINDLE ORGANIZATION | 5.99E-07 | -2.1570196 | 118 |
| GOBP INTERLEUKIN 1 BETA PRODUCTION | 6.12E-07 | 2.04888157 | 83 |
| GOBP FC RECEPTOR SIGNALING PATHWAY | 6.35E-07 | 1.90279506 | 170 |
| GOBP NEUTROPHIL MIGRATION | 6.63E-07 | 1.99656393 | 111 |
| GOBP TRNA METABOLIC PROCESS | 7.19E-07 | -2.0692232 | 178 |
| GOBP REGULATION OF LEUKOCYTE MIGRATION | 7.25E-07 | 1.88521012 | 183 |
| GOBP DNA REPLICATION INITIATION | 7.28E-07 | -2.4233686 | 39 |
| GOBP ANTIGEN PROCESSING AND PRESENTATION OF PEPTIDE ANTIGEN | 7.44E-07 | 1.87910985 | 185 |
| GOBP INTERLEUKIN 6 PRODUCTION | 7.44E-07 | 1.94770762 | 127 |
| GOBP INTERLEUKIN 2 PRODUCTION | 8.04E-07 | 2.09359789 | 56 |
| GOBP REGULATION OF ALPHA BETA T CELL ACTIVATION | 8.49E-07 | 2.03348952 | 91 |
| GOBP CHEMICAL HOMEOSTASIS | 1.09E-06 | 1.4479912 | 985 |
| GOBP CHROMOSOME LOCALIZATION | 1.09E-06 | -2.2943324 | 76 |
| GOBP GRANULOCYTE CHEMOTAXIS | 1.09E-06 | 1.96399486 | 114 |
| GOBP POSITIVE REGULATION OF TRANSPORT | 1.09E-06 | 1.5051944 | 780 |
| GOBP POSITIVE REGULATION OF TUMOR NECROSIS FACTOR SUPERFAMILY CYTOKINE PRODUCTION | 1.09E-06 | 2.02249735 | 80 |
| GOBP CATION TRANSPORT | 1.15E-06 | 1.46803404 | 950 |
| GOBP MONOCYTE CHEMOTAXIS | 1.16E-06 | 2.11580949 | 57 |
| GOBP NEGATIVE REGULATION OF MITOTIC CELL CYCLE | 1.37E-06 | -1.8179423 | 288 |
| GOBP REGULATION OF B CELL PROLIFERATION | 1.41E-06 | 2.10207747 | 53 |

| | | | |
|---|---|---|---|
| GOBP INTERFERON GAMMA MEDIATED SIGNALING PATHWAY | 1.52E-06 | 2.01617782 | 85 |
| GOBP METAPHASE PLATE CONGRESSION | 1.56E-06 | -2.3011843 | 63 |
| GOBP DNA BIOSYNTHETIC PROCESS | 1.62E-06 | -2.0379286 | 180 |
| GOBP COMPLEMENT ACTIVATION | 1.83E-06 | 2.11020048 | 50 |
| GOBP CD4 POSITIVE ALPHA BETA T CELL DIFFERENTIATION | 1.97E-06 | 2.04214464 | 72 |
| GOBP NEGATIVE REGULATION OF RESPONSE TO EXTERNAL STIMULUS | 1.97E-06 | 1.70191504 | 315 |
| GOBP DNA PACKAGING | 2.09E-06 | -1.9776102 | 190 |
| GOBP LYMPHOCYTE CHEMOTAXIS | 2.16E-06 | 2.10387514 | 50 |
| GOBP CD4 POSITIVE ALPHA BETA T CELL ACTIVATION | 2.43E-06 | 1.97549679 | 89 |
| GOBP MYELOID LEUKOCYTE DIFFERENTIATION | 2.70E-06 | 1.86312533 | 181 |
| GOBP CYTOPLASMIC TRANSLATION | 2.75E-06 | -2.1510905 | 98 |
| GOBP POSITIVE REGULATION OF PHAGOCYTOSIS | 2.75E-06 | 2.09593276 | 57 |
| GOBP TRANSLATIONAL ELONGATION | 2.98E-06 | -2.0677531 | 131 |
| GOBP RIBOSOME ASSEMBLY | 3.29E-06 | -2.2845639 | 59 |
| GOBP RNA SPLICING VIA TRANSESTERIFICATION REACTIONS | 3.79E-06 | -1.68959 | 338 |
| GOBP PROTEIN LOCALIZATION TO CHROMOSOME | 4.00E-06 | -2.1580645 | 89 |
| GOBP B CELL MEDIATED IMMUNITY | 4.04E-06 | 1.96587784 | 106 |
| GOBP DNA INTEGRITY CHECKPOINT | 4.10E-06 | -1.9817434 | 153 |
| GOBP NEUTROPHIL CHEMOTAXIS | 4.23E-06 | 1.96645406 | 94 |
| GOBP CENTROMERE COMPLEX ASSEMBLY | 4.36E-06 | -2.308137 | 50 |
| GOBP MICROGLIAL CELL ACTIVATION | 4.53E-06 | 2.0637595 | 42 |

| | | | |
|---|---|---|---|
| GOBP EMBRYONIC SKELETAL SYSTEM DEVELOPMENT | 4.53E-06 | -2.0843672 | 106 |
| GOBP NEGATIVE REGULATION OF NUCLEAR DIVISION | 4.87E-06 | -2.23751 | 51 |
| GOBP POSITIVE REGULATION OF GENE EXPRESSION | 5.01E-06 | 1.43571375 | 937 |
| GOBP OSTEOCLAST DIFFERENTIATION | 5.11E-06 | 1.98202008 | 79 |
| GOBP REGULATION OF DNA REPLICATION | 5.11E-06 | -2.1445915 | 101 |
| GOBP PATTERN SPECIFICATION PROCESS | 5.17E-06 | -1.6976804 | 347 |
| GOBP REGULATION OF PATTERN RECOGNITION RECEPTOR SIGNALING PATHWAY | 5.36E-06 | 1.95738319 | 94 |
| GOBP DENDRITIC CELL DIFFERENTIATION | 5.53E-06 | 2.04898373 | 43 |
| GOBP RESPONSE TO MOLECULE OF BACTERIAL ORIGIN | 5.67E-06 | 1.69654671 | 303 |
| GOBP TELOMERE ORGANIZATION | 5.67E-06 | -1.9525866 | 160 |
| GOBP NEGATIVE REGULATION OF LEUKOCYTE MEDIATED IMMUNITY | 6.17E-06 | 2.05307993 | 49 |
| GOBP NEGATIVE REGULATION OF LYMPHOCYTE MEDIATED IMMUNITY | 6.27E-06 | 2.05522575 | 42 |
| GOBP REGULATION OF ANTIGEN PROCESSING AND PRESENTATION | 6.53E-06 | 2.06172943 | 20 |
| GOBP REGULATION OF T CELL MEDIATED IMMUNITY | 6.53E-06 | 2.01357329 | 70 |
| GOBP COLLAGEN FIBRIL ORGANIZATION | 6.80E-06 | -2.2847992 | 50 |
| GOBP REGULATION OF TOLL LIKE RECEPTOR SIGNALING PATHWAY | 6.85E-06 | 2.02350966 | 68 |
| GOBP CALCIUM ION TRANSPORT | 6.91E-06 | 1.67943515 | 332 |
| GOBP PRODUCTION OF MOLECULAR MEDIATOR OF IMMUNE RESPONSE | 7.31E-06 | 1.83056494 | 170 |

| | | | |
|---|---|---|---|
| GOBP REGULATION OF B CELL RECEPTOR SIGNALING PATHWAY | 7.48E-06 | 2.06607268 | 24 |
| GOBP CELLULAR RESPONSE TO BIOTIC STIMULUS | 7.76E-06 | 1.79352392 | 207 |
| GOBP REGULATION OF PRODUCTION OF MOLECULAR MEDIATOR OF IMMUNE RESPONSE | 7.76E-06 | 1.90407059 | 131 |
| GOBP REGULATION OF COMPLEMENT ACTIVATION | 8.22E-06 | 2.06354398 | 40 |
| GOBP NEGATIVE REGULATION OF DEFENSE RESPONSE | 8.93E-06 | 1.81553887 | 176 |
| GOBP RECEPTOR MEDIATED ENDOCYTOSIS | 1.17E-05 | 1.75651463 | 234 |
| GOBP DNA RECOMBINATION | 1.25E-05 | -1.7485396 | 274 |
| GOBP POSITIVE REGULATION OF GTPASE ACTIVITY | 1.30E-05 | 1.60063502 | 388 |
| GOBP PROTEIN LOCALIZATION TO CHROMOSOME CENTROMERIC REGION | 1.56E-05 | -2.2854634 | 25 |
| GOBP RESPONSE TO CHEMOKINE | 1.56E-05 | 1.94431265 | 84 |
| GOBP ANTERIOR POSTERIOR PATTERN SPECIFICATION | 1.61E-05 | -1.9053267 | 164 |
| GOBP NEGATIVE REGULATION OF ADAPTIVE IMMUNE RESPONSE | 1.68E-05 | 2.04546845 | 48 |
| GOBP REGULATION OF GTPASE ACTIVITY | 2.12E-05 | 1.54149285 | 460 |
| GOBP T CELL MEDIATED IMMUNITY | 2.18E-05 | 1.93054271 | 92 |
| GOBP EMBRYONIC ORGAN MORPHOGENESIS | 2.19E-05 | -1.7816555 | 243 |
| GOBP CELLULAR DEFENSE RESPONSE | 2.22E-05 | 2.04476554 | 41 |
| GOBP CELLULAR HOMEOSTASIS | 2.22E-05 | 1.44521791 | 795 |

| | | | |
|---|---|---|---|
| GOBP MITOTIC METAPHASE PLATE CONGRESSION | 2.22E-05 | -2.2255679 | 50 |
| GOBP MACROPHAGE MIGRATION | 2.25E-05 | 2.0293011 | 50 |
| GOBP POSITIVE REGULATION OF LEUKOCYTE MIGRATION | 2.29E-05 | 1.86645957 | 122 |
| GOBP PROTEIN DNA COMPLEX SUBUNIT ORGANIZATION | 2.39E-05 | -1.817709 | 234 |
| GOBP CYTOKINE PRODUCTION INVOLVED IN IMMUNE RESPONSE | 2.48E-05 | 1.93438574 | 86 |
| GOBP MYELOID CELL DIFFERENTIATION | 2.51E-05 | 1.62901757 | 372 |
| GOBP REGULATION OF MYELOID LEUKOCYTE MEDIATED IMMUNITY | 2.51E-05 | 2.02520781 | 50 |
| GOBP NEGATIVE REGULATION OF LEUKOCYTE APOPTOTIC PROCESS | 2.67E-05 | 2.00573844 | 44 |
| GOBP REGULATION OF CHEMOTAXIS | 2.67E-05 | 1.78253714 | 198 |
| GOBP T CELL RECEPTOR SIGNALING PATHWAY | 2.72E-05 | 1.79918069 | 187 |
| GOBP ATTACHMENT OF SPINDLE MICROTUBULES TO KINETOCHORE | 2.97E-05 | -2.2807739 | 35 |
| GOBP CELLULAR RESPONSE TO MOLECULE OF BACTERIAL ORIGIN | 3.05E-05 | 1.77809828 | 183 |
| GOBP MATURATION OF SSU RRNA | 3.06E-05 | -2.2176035 | 48 |
| GOBP ENDOCARDIAL CUSHION DEVELOPMENT | 3.10E-05 | -2.2472121 | 39 |
| GOBP SUPEROXIDE METABOLIC PROCESS | 3.17E-05 | 1.98987591 | 58 |
| GOBP MEIOSIS I CELL CYCLE PROCESS | 3.28E-05 | -1.9874971 | 102 |
| GOBP NEGATIVE REGULATION OF T CELL MEDIATED IMMUNITY | 3.28E-05 | 2.0250169 | 20 |
| GOBP REGULATORY T CELL DIFFERENTIATION | 3.39E-05 | 2.02695015 | 32 |
| GOBP REGULATION OF REGULATED SECRETORY PATHWAY | 3.60E-05 | 1.82882611 | 128 |

| | | | |
|---|---|---|---|
| GOBP POSITIVE REGULATION OF MYELOID LEUKOCYTE MEDIATED IMMUNITY | 3.66E-05 | 2.03235254 | 17 |
| GOBP MESENCHYMAL CELL DIFFERENTIATION | 3.91E-05 | -1.843969 | 206 |
| GOBP EMBRYONIC SKELETAL SYSTEM MORPHOGENESIS | 3.94E-05 | -2.0617286 | 82 |
| GOBP NEGATIVE REGULATION OF INTERLEUKIN 12 PRODUCTION | 3.98E-05 | 2.02944723 | 17 |
| GOBP COMPLEMENT RECEPTOR MEDIATED SIGNALING PATHWAY | 4.15E-05 | 1.92200733 | 10 |
| GOBP POSITIVE REGULATION OF CHEMOTAXIS | 4.34E-05 | 1.82668764 | 127 |
| GOBP REGULATION OF LEUKOCYTE CHEMOTAXIS | 4.34E-05 | 1.89093024 | 106 |
| GOBP DNA REPLICATION INDEPENDENT NUCLEOSOME ORGANIZATION | 4.37E-05 | -2.2026057 | 48 |
| GOBP DNA DEPENDENT DNA REPLICATION MAINTENANCE OF FIDELITY | 4.40E-05 | -2.2050475 | 47 |
| GOBP REGULATION OF MACROPHAGE MIGRATION | 4.57E-05 | 2.02737594 | 38 |
| GOBP CALCIUM MEDIATED SIGNALING | 4.58E-05 | 1.76253943 | 169 |
| GOBP POSITIVE REGULATION OF MACROPHAGE MIGRATION | 4.58E-05 | 2.01768845 | 24 |
| GOBP RNA LOCALIZATION | 4.84E-05 | -1.7787864 | 221 |
| GOBP NEGATIVE REGULATION OF B CELL ACTIVATION | 4.84E-05 | 2.01594312 | 32 |
| GOBP KINETOCHORE ORGANIZATION | 5.62E-05 | -2.2410265 | 22 |
| GOBP POSITIVE REGULATION OF ALPHA BETA T CELL ACTIVATION | 5.62E-05 | 1.99171376 | 59 |

| | | | |
|---|---|---|---|
| GOBP UROGENITAL SYSTEM DEVELOPMENT | 5.96E-05 | -1.6111749 | 290 |
| GOBP NEGATIVE REGULATION OF RESPONSE TO BIOTIC STIMULUS | 5.99E-05 | 1.85966297 | 89 |
| GOBP REGULATION OF MONONUCLEAR CELL MIGRATION | 6.01E-05 | 1.86970798 | 99 |
| GOBP HUMORAL IMMUNE RESPONSE MEDIATED BY CIRCULATING IMMUNOGLOBULIN | 6.36E-05 | 1.97467049 | 42 |
| GOBP REGULATION OF MACROPHAGE ACTIVATION | 6.46E-05 | 1.98081879 | 52 |
| GOBP CONNECTIVE TISSUE DEVELOPMENT | 6.64E-05 | -1.818719 | 214 |
| GOBP EXTERNAL ENCAPSULATING STRUCTURE ORGANIZATION | 6.64E-05 | -1.5146238 | 357 |
| GOBP CARDIAC CHAMBER DEVELOPMENT | 6.98E-05 | -1.8793643 | 137 |
| GOBP ANTIGEN PROCESSING AND PRESENTATION OF PEPTIDE OR POLYSACCHARIDE ANTIGEN VIA MHC CLASS II | 7.13E-05 | 1.8538163 | 100 |
| GOBP ACTIVATION OF INNATE IMMUNE RESPONSE | 7.24E-05 | 1.79798664 | 136 |
| GOBP BRANCHING MORPHOGENESIS OF AN EPITHELIAL TUBE | 0.00010335 | -1.8648983 | 128 |
| GOBP CHROMATIN REMODELING AT CENTROMERE | 0.00010767 | -2.2213063 | 41 |
| GOBP RESPONSE TO TUMOR NECROSIS FACTOR | 0.00011075 | 1.62898557 | 284 |
| GOBP PRODUCTION OF MOLECULAR MEDIATOR INVOLVED IN INFLAMMATORY RESPONSE | 0.00011845 | 1.96312297 | 61 |

| | | | |
|---|---|---|---|
| GOBP MESENCHYME MORPHOGENESIS | 0.00012226 | -2.2245863 | 43 |
| GOBP REGULATION OF CYTOSOLIC CALCIUM ION CONCENTRATION | 0.00012729 | 1.62628469 | 266 |
| GOBP REGULATION OF DNA DEPENDENT DNA REPLICATION | 0.0001298 | -2.1555938 | 46 |
| GOBP REGULATION OF CELL DIVISION | 0.00013523 | -1.807188 | 148 |
| GOBP HISTONE EXCHANGE | 0.00013988 | -2.1362236 | 49 |
| GOBP T CELL MEDIATED CYTOTOXICITY | 0.00014108 | 1.97997309 | 41 |
| GOBP T CELL DIFFERENTIATION INVOLVED IN IMMUNE RESPONSE | 0.00014543 | 1.92415086 | 64 |
| GOBP INNATE IMMUNE RESPONSE ACTIVATING SIGNAL TRANSDUCTION | 0.00015049 | 1.82164664 | 109 |
| GOBP REGIONALIZATION | 0.00015127 | -1.6568691 | 269 |
| GOBP REGULATION OF B CELL MEDIATED IMMUNITY | 0.00015607 | 1.92875967 | 51 |
| GOBP CHEMOKINE PRODUCTION | 0.00015862 | 1.86309261 | 79 |
| GOBP NEGATIVE REGULATION OF B CELL PROLIFERATION | 0.00016253 | 1.97480043 | 16 |
| GOBP NEGATIVE REGULATION OF TUMOR NECROSIS FACTOR SUPERFAMILY CYTOKINE PRODUCTION | 0.00016386 | 1.92526573 | 51 |
| GOBP SECOND MESSENGER MEDIATED SIGNALING | 0.00016502 | 1.63418239 | 251 |
| GOBP B CELL DIFFERENTIATION | 0.00016663 | 1.80358578 | 115 |
| GOBP MATURATION OF 5 8S RRNA | 0.00016663 | -2.1836055 | 35 |
| GOBP MITOTIC DNA INTEGRITY CHECKPOINT | 0.00016663 | -1.9043318 | 104 |
| GOBP LYMPHOCYTE COSTIMULATION | 0.00016678 | 1.93985298 | 53 |
| GOBP REGULATION OF SECRETION | 0.00016781 | 1.48388172 | 523 |

| | | | |
|---|---|---|---|
| GOBP NEGATIVE REGULATION OF HEMOPOIESIS | 0.0001734 | 1.83415789 | 93 |
| GOBP REGULATION OF ALPHA BETA T CELL DIFFERENTIATION | 0.00018137 | 1.92718716 | 60 |
| GOBP IMMUNE RESPONSE INHIBITING SIGNAL TRANSDUCTION | 0.00018148 | 1.80566238 | 7 |
| GOBP DENDRITIC CELL CHEMOTAXIS | 0.00018926 | 1.9657981 | 22 |
| GOBP POSITIVE REGULATION OF ALPHA BETA T CELL DIFFERENTIATION | 0.00018926 | 1.94845983 | 45 |
| GOBP LIPOPOLYSACCHARIDE MEDIATED SIGNALING PATHWAY | 0.00019545 | 1.94640676 | 55 |
| GOBP SPROUTING ANGIOGENESIS | 0.00019545 | -1.8945511 | 115 |
| GOBP LEUKOCYTE APOPTOTIC PROCESS | 0.00019726 | 1.84563385 | 95 |
| GOBP REGULATION OF INFLAMMATORY RESPONSE TO ANTIGENIC STIMULUS | 0.00019726 | 1.96292501 | 25 |
| GOBP CELL MIGRATION INVOLVED IN SPROUTING ANGIOGENESIS | 0.00020043 | -2.0345552 | 51 |
| GOBP INTERSTRAND CROSS LINK REPAIR | 0.00020076 | -2.0868613 | 57 |
| GOBP NEGATIVE REGULATION OF TOLL LIKE RECEPTOR SIGNALING PATHWAY | 0.00020076 | 1.97737714 | 38 |
| GOBP ANAPHASE PROMOTING COMPLEX DEPENDENT CATABOLIC PROCESS | 0.00023063 | -2.0121413 | 80 |
| GOBP NATURAL KILLER CELL MEDIATED IMMUNITY | 0.00023063 | 1.93103059 | 53 |
| GOBP REGULATION OF DNA BIOSYNTHETIC PROCESS | 0.00023509 | -1.8820608 | 104 |
| GOBP TRNA PROCESSING | 0.00023509 | -1.848264 | 129 |

| | | | |
|---|---|---|---|
| GOBP REGULATION OF LEUKOCYTE APOPTOTIC PROCESS | 0.00023516 | 1.87281377 | 75 |
| GOBP INTERLEUKIN 10 PRODUCTION | 0.00023713 | 1.91322007 | 51 |
| GOBP MATURATION OF LSU RRNA | 0.00024764 | -2.2023809 | 28 |
| GOBP REGULATION OF EXOCYTOSIS | 0.00024792 | 1.71609067 | 186 |
| GOBP MITOTIC SPINDLE ASSEMBLY | 0.00025716 | -2.0015692 | 65 |
| GOBP POSITIVE REGULATION OF LEUKOCYTE CHEMOTAXIS | 0.00025993 | 1.85494957 | 81 |
| GOBP MAST CELL ACTIVATION | 0.00026058 | 1.94859068 | 57 |
| GOBP SPLICEOSOMAL SNRNP ASSEMBLY | 0.00026196 | -2.1417817 | 39 |
| GOBP CARDIAC VENTRICLE DEVELOPMENT | 0.00026207 | -1.9498551 | 103 |
| GOBP DENDRITIC CELL MIGRATION | 0.00026807 | 1.94964572 | 27 |
| GOBP ATTACHMENT OF MITOTIC SPINDLE MICROTUBULES TO KINETOCHORE | 0.00027115 | -2.1520742 | 15 |
| GOBP DNA STRAND ELONGATION | 0.00028466 | -2.1332393 | 25 |
| GOBP RIBOSOMAL LARGE SUBUNIT ASSEMBLY | 0.00028516 | -2.1979032 | 26 |
| GOBP NEGATIVE REGULATION OF MULTICELLULAR ORGANISMAL PROCESS | 0.00029614 | 1.38162363 | 863 |
| GOBP CELL JUNCTION ASSEMBLY | 0.00031128 | -1.4581754 | 383 |
| GOBP REGULATION OF CD4 POSITIVE ALPHA BETA T CELL DIFFERENTIATION | 0.00031128 | 1.92932547 | 45 |
| GOBP VIRAL GENE EXPRESSION | 0.00032703 | -1.7430349 | 198 |
| GOBP REGULATION OF LEUKOCYTE DEGRANULATION | 0.00034137 | 1.92187023 | 40 |
| GOBP DEFENSE RESPONSE TO VIRUS | 0.00034668 | 1.63632652 | 227 |

| | | | |
|---|---|---|---|
| GOBP REPLICATION FORK PROCESSING | 0.00035136 | -2.1342906 | 38 |
| GOBP POSITIVE REGULATION OF INTERLEUKIN 12 PRODUCTION | 0.00035328 | 1.95149273 | 34 |
| GOBP PURINERGIC NUCLEOTIDE RECEPTOR SIGNALING PATHWAY | 0.00036437 | 1.93646391 | 27 |
| GOBP MACROPHAGE CHEMOTAXIS | 0.00037942 | 1.93680509 | 37 |
| GOBP RESPONSE TO TYPE I INTERFERON | 0.0003857 | 1.78211701 | 89 |
| GOBP HEART MORPHOGENESIS | 0.0003868 | -1.6615214 | 211 |
| GOBP POSITIVE REGULATION OF INTERLEUKIN 1 PRODUCTION | 0.0003868 | 1.92290396 | 55 |
| GOBP MEIOTIC CHROMOSOME SEGREGATION | 0.00038726 | -2.0008489 | 72 |
| GOBP POSITIVE REGULATION OF PHOSPHATASE ACTIVITY | 0.00038726 | 1.91705653 | 30 |
| GOBP PROTEIN LOCALIZATION TO KINETOCHORE | 0.00039369 | -2.1348705 | 19 |
| GOBP POSITIVE REGULATION OF INTERLEUKIN 2 PRODUCTION | 0.0003959 | 1.94846363 | 29 |
| GOBP NEGATIVE REGULATION OF CELL CYCLE G2 M PHASE TRANSITION | 0.00039595 | -1.8236645 | 105 |
| GOBP POSITIVE REGULATION OF INFLAMMATORY RESPONSE | 0.00041248 | 1.75676751 | 112 |
| GOBP POSITIVE REGULATION OF DNA BIOSYNTHETIC PROCESS | 0.00043595 | -1.9617357 | 65 |
| GOBP REGULATION OF PLASMA LIPOPROTEIN PARTICLE LEVELS | 0.00046131 | 1.84382232 | 75 |
| GOBP POSITIVE REGULATION OF ADAPTIVE IMMUNE RESPONSE | 0.00047459 | 1.8116441 | 95 |

| | | | |
|---|---|---|---|
| GOBP POSITIVE REGULATION OF PRODUCTION OF MOLECULAR MEDIATOR OF IMMUNE RESPONSE | 0.0004852 | 1.77869248 | 90 |
| GOBP POSITIVE REGULATION OF ANTIGEN RECEPTOR MEDIATED SIGNALING PATHWAY | 0.000498 | 1.92265974 | 21 |
| GOBP SIGNAL TRANSDUCTION BY P53 CLASS MEDIATOR | 0.0005077 | -1.5811679 | 255 |
| GOBP MATURATION OF SSU RRNA FROM TRICISTRONIC RRNA TRANSCRIPT SSU RRNA 5 8S RRNA LSU RRNA | 0.00054035 | -2.0965897 | 35 |
| GOBP RESPONSE TO VIRUS | 0.00054356 | 1.55277239 | 316 |
| GOBP POSITIVE REGULATION OF INTERLEUKIN 6 PRODUCTION | 0.00055029 | 1.83573848 | 77 |
| GOBP CD8 POSITIVE ALPHA BETA T CELL ACTIVATION | 0.00056369 | 1.95105505 | 23 |
| GOBP EPITHELIAL TO MESENCHYMAL TRANSITION | 0.00057278 | -1.7654006 | 136 |
| GOBP POSITIVE REGULATION OF CD4 POSITIVE ALPHA BETA T CELL DIFFERENTIATION | 0.00059235 | 1.92143334 | 28 |
| GOBP SKELETAL SYSTEM MORPHOGENESIS | 0.00059789 | -1.6947936 | 190 |
| GOBP RUFFLE ORGANIZATION | 0.00060261 | 1.89563769 | 54 |
| GOBP POSITIVE REGULATION OF CYTOKINE PRODUCTION INVOLVED IN IMMUNE RESPONSE | 0.00060359 | 1.88267254 | 50 |
| GOBP SYNAPSE PRUNING | 0.00060359 | 1.86043033 | 11 |
| GOBP DNA TEMPLATED TRANSCRIPTION TERMINATION | 0.00062538 | -1.9886717 | 75 |
| GOBP FEMALE MEIOTIC NUCLEAR DIVISION | 0.00064699 | -2.1045468 | 23 |

| | | | |
|---|---|---|---|
| GOBP SUPEROXIDE ANION GENERATION | 0.00064708 | 1.89131613 | 30 |
| GOBP REACTIVE OXYGEN SPECIES METABOLIC PROCESS | 0.00066422 | 1.60149359 | 237 |
| GOBP SENSORY ORGAN MORPHOGENESIS | 0.00066422 | -1.6911492 | 202 |
| GOBP HOMOLOGOUS RECOMBINATION | 0.00067439 | -1.9655039 | 51 |
| GOBP INFLAMMATORY RESPONSE TO ANTIGENIC STIMULUS | 0.00074522 | 1.86558948 | 51 |
| GOBP DNA STRAND ELONGATION INVOLVED IN DNA REPLICATION | 0.00076013 | -2.1034378 | 19 |
| GOBP ORGANIC HYDROXY COMPOUND TRANSPORT | 0.00076013 | 1.6202863 | 208 |
| GOBP REGULATION OF PHOSPHATIDYLINOSITOL 3 KINASE ACTIVITY | 0.00077339 | 1.88039451 | 54 |
| GOBP IMMUNOLOGICAL SYNAPSE FORMATION | 0.0007779 | 1.87354111 | 13 |
| GOBP SPINDLE ASSEMBLY | 0.00078411 | -1.8630085 | 112 |
| GOBP CARTILAGE DEVELOPMENT | 0.00079785 | -1.6955655 | 161 |
| GOBP REGULATION OF T CELL MEDIATED CYTOTOXICITY | 0.00080699 | 1.90703535 | 33 |
| GOBP POSITIVE REGULATION OF CD8 POSITIVE ALPHA BETA T CELL DIFFERENTIATION | 0.00080914 | 1.70146983 | 5 |
| GOBP REGULATION OF NATURAL KILLER CELL MEDIATED IMMUNITY | 0.00080914 | 1.89676271 | 37 |
| GOBP REGULATION OF SYNCYTIUM FORMATION BY PLASMA MEMBRANE FUSION | 0.00086507 | 1.90989449 | 22 |
| GOBP APPENDAGE DEVELOPMENT | 0.00089008 | -1.7008781 | 148 |
| GOBP EOSINOPHIL MIGRATION | 0.00091104 | 1.92296363 | 19 |
| GOBP REGULATION OF MYELOID LEUKOCYTE DIFFERENTIATION | 0.00094596 | 1.75978707 | 99 |

Table S7. GSEA of KEGG pathways of endothelial cells in infected chips compared to mock infected chips, both with macrophages (adj $P < 0.05$).

| **Gene set** | **Adjusted *P*-value** | **NES** | **Size** |
|---|---|---|---|
| KEGG LEISHMANIA INFECTION | 0.000009 | -2.00011 | 66 |
| KEGG CYTOSOLIC DNA SENSING PATHWAY | 0.000085 | 2.163961 | 42 |
| KEGG AMINO SUGAR AND NUCLEOTIDE SUGAR METABOLISM | 0.000536 | -1.91461 | 44 |
| KEGG RIG I LIKE RECEPTOR SIGNALING PATHWAY | 0.001519 | 2.045047 | 55 |
| KEGG LEUKOCYTE TRANSENDOTHELIAL MIGRATION | 0.001987 | -1.75706 | 104 |
| KEGG ASTHMA | 0.002178 | -1.84934 | 19 |
| KEGG FC EPSILON RI SIGNALING PATHWAY | 0.002178 | -1.79255 | 68 |
| KEGG FC GAMMA R MEDIATED PHAGOCYTOSIS | 0.002178 | -1.73245 | 93 |
| KEGG LYSOSOME | 0.002178 | -1.68343 | 119 |
| KEGG HEMATOPOIETIC CELL LINEAGE | 0.003463 | -1.73361 | 73 |
| KEGG B CELL RECEPTOR SIGNALING PATHWAY | 0.009915 | -1.68093 | 73 |
| KEGG CELL ADHESION MOLECULES CAMS | 0.010454 | -1.62085 | 124 |
| KEGG INTESTINAL IMMUNE NETWORK FOR IGA PRODUCTION | 0.023397 | -1.71709 | 37 |
| KEGG RIBOSOME | 0.040085 | -1.55295 | 85 |
| KEGG COMPLEMENT AND COAGULATION CASCADES | 0.044648 | -1.59412 | 58 |

Table S8. GSEA of HALLMARK pathways of endothelial cells in infected chips compared to mock infected chips, both with macrophages (adj $P < 0.05$).

| **Gene set** | **Adjusted *P*-value** | **NES** | **Size** |
|---|---|---|---|
| HALLMARK INTERFERON ALPHA RESPONSE | 1.936968E-25 | 2.807312 | 97 |
| HALLMARK INTERFERON GAMMA RESPONSE | 4.011598E-21 | 2.533471 | 197 |
| HALLMARK COAGULATION | 0.001057 | -1.66203 | 118 |
| HALLMARK ESTROGEN RESPONSE LATE | 0.001848 | -1.61101 | 193 |
| HALLMARK ESTROGEN RESPONSE EARLY | 0.010603 | -1.50308 | 198 |
| HALLMARK ALLOGRAFT REJECTION | 0.020331 | -1.47559 | 183 |
| HALLMARK KRAS SIGNALING UP | 0.030545 | -1.40512 | 190 |
| HALLMARK XENOBIOTIC METABOLISM | 0.030545 | -1.45735 | 175 |
| HALLMARK COMPLEMENT | 0.038247 | -1.39645 | 188 |
| HALLMARK BILE ACID METABOLISM | 0.047161 | -1.46773 | 96 |

Table S9. GOBP analysis of endothelial cells in infected chips compared to mock infected chips, both with macrophages (adj $P < 0.001$).

| **Gene set** | **Adjusted *P*-value** | **NES** | **Size** |
|---|---|---|---|
| GOBP DEFENSE RESPONSE TO VIRUS | 9.44E-21 | 2.55883831 | 222 |
| GOBP MYELOID LEUKOCYTE ACTIVATION | 5.75E-18 | -1.8843586 | 602 |
| GOBP RESPONSE TO VIRUS | 1.00E-16 | 2.30020015 | 307 |
| GOBP CELL ACTIVATION INVOLVED IN IMMUNE RESPONSE | 5.94E-16 | -1.8265824 | 644 |
| GOBP LEUKOCYTE MEDIATED IMMUNITY | 1.73E-15 | -1.8012391 | 690 |
| GOBP VIRAL GENOME REPLICATION | 1.73E-15 | 2.55535397 | 122 |
| GOBP RESPONSE TO TYPE I INTERFERON | 1.21E-14 | 2.6343501 | 85 |
| GOBP MYELOID LEUKOCYTE MEDIATED IMMUNITY | 1.29E-14 | -1.8710855 | 505 |
| GOBP EXOCYTOSIS | 3.80E-14 | -1.7066099 | 811 |
| GOBP REGULATION OF VIRAL GENOME REPLICATION | 3.28E-13 | 2.62248685 | 79 |
| GOBP NEGATIVE REGULATION OF VIRAL PROCESS | 5.32E-13 | 2.6147497 | 79 |
| GOBP REGULATION OF BIOLOGICAL PROCESS INVOLVED IN SYMBIOTIC INTERACTION | 2.83E-12 | 2.30987452 | 181 |
| GOBP NEGATIVE REGULATION OF VIRAL GENOME REPLICATION | 1.54E-11 | 2.53155684 | 51 |
| GOBP REGULATION OF VIRAL LIFE CYCLE | 5.65E-11 | 2.40516526 | 134 |
| GOBP CELL CELL ADHESION | 4.63E-09 | -1.60135 | 771 |
| GOBP PHAGOCYTOSIS | 3.58E-08 | -1.8907881 | 249 |
| GOBP ORGANIC ACID METABOLIC PROCESS | 3.93E-08 | -1.5435402 | 900 |
| GOBP REGULATION OF CELL ACTIVATION | 3.93E-08 | -1.6841469 | 503 |

| | | | |
|---|---|---|---|
| GOBP ORGANIC HYDROXY COMPOUND METABOLIC PROCESS | 6.27E-07 | -1.6735028 | 457 |
| GOBP MONOCARBOXYLIC ACID METABOLIC PROCESS | 2.01E-06 | -1.6162068 | 532 |
| GOBP CELLULAR LIPID METABOLIC PROCESS | 2.99E-06 | -1.4817997 | 897 |
| GOBP CELLULAR RESPONSE TO DSRNA | 2.99E-06 | 2.23884753 | 21 |
| GOBP IMMUNE RESPONSE REGULATING SIGNALING PATHWAY | 3.89E-06 | -1.7033541 | 365 |
| GOBP LEUKOCYTE DIFFERENTIATION | 6.95E-06 | -1.631157 | 461 |
| GOBP REGULATION OF VESICLE MEDIATED TRANSPORT | 8.05E-06 | -1.618595 | 459 |
| GOBP ORGANIC HYDROXY COMPOUND TRANSPORT | 8.47E-06 | -1.8090194 | 198 |
| GOBP POSITIVE REGULATION OF CELL ACTIVATION | 1.29E-05 | -1.7171392 | 308 |
| GOBP LYMPHOCYTE ACTIVATION | 1.61E-05 | -1.5583722 | 600 |
| GOBP RESPONSE TO INTERFERON BETA | 1.95E-05 | 2.22630172 | 27 |
| GOBP VIRAL LIFE CYCLE | 2.22E-05 | 1.71531617 | 328 |
| GOBP CELLULAR RESPONSE TO EXOGENOUS DSRNA | 2.67E-05 | 2.17209668 | 17 |
| GOBP MACROPHAGE ACTIVATION | 2.75E-05 | -1.9187557 | 91 |
| GOBP MEMBRANE INVAGINATION | 2.75E-05 | -1.9547349 | 69 |
| GOBP OSTEOCLAST DIFFERENTIATION | 2.99E-05 | -1.9413455 | 81 |
| GOBP CATION TRANSPORT | 3.19E-05 | -1.4337874 | 904 |
| GOBP LEUKOCYTE PROLIFERATION | 3.86E-05 | -1.7163164 | 264 |
| GOBP POSITIVE REGULATION OF TRANSPORT | 4.25E-05 | -1.467468 | 772 |
| GOBP REGULATION OF IMMUNE RESPONSE | 4.29E-05 | -1.4732429 | 758 |
| GOBP REGULATION OF HORMONE LEVELS | 5.24E-05 | -1.6201569 | 409 |

| | | | |
|---|---|---|---|
| GOBP SMALL MOLECULE BIOSYNTHETIC PROCESS | 5.24E-05 | -1.5150359 | 595 |
| GOBP CELL CELL ADHESION VIA PLASMA MEMBRANE ADHESION MOLECULES | 5.46E-05 | -1.7231079 | 233 |
| GOBP INFLAMMATORY RESPONSE | 5.55E-05 | -1.5022814 | 612 |
| GOBP ORGANIC ACID BIOSYNTHETIC PROCESS | 5.83E-05 | -1.701087 | 277 |
| GOBP AMYLOID BETA CLEARANCE | 7.14E-05 | -1.9652764 | 35 |
| GOBP LEUKOCYTE MIGRATION | 8.33E-05 | -1.6180122 | 383 |
| GOBP RESPONSE TO INTERFERON ALPHA | 9.05E-05 | 2.14056909 | 21 |
| GOBP REGULATION OF B CELL ACTIVATION | 9.47E-05 | -1.8501419 | 113 |
| GOBP REGULATION OF SECRETION | 0.00011652 | -1.5484011 | 502 |
| GOBP ADAPTIVE IMMUNE RESPONSE | 0.00012225 | -1.6222256 | 353 |
| GOBP MACROPHAGE FUSION | 0.00013012 | -1.7213119 | 6 |
| GOBP ANTIGEN PROCESSING AND PRESENTATION OF PEPTIDE OR POLYSACCHARIDE ANTIGEN VIA MHC CLASS II | 0.00015532 | -1.8642131 | 100 |
| GOBP MULTICELLULAR ORGANISMAL HOMEOSTASIS | 0.0001988 | -1.558721 | 440 |
| GOBP MAPK CASCADE | 0.00021156 | -1.4243207 | 788 |
| GOBP REGULATION OF LYMPHOCYTE ACTIVATION | 0.00026693 | -1.5761441 | 395 |
| GOBP POSITIVE REGULATION OF IMMUNE SYSTEM PROCESS | 0.00028834 | -1.4166316 | 827 |
| GOBP REGULATION OF LEUKOCYTE PROLIFERATION | 0.00029998 | -1.7153016 | 210 |
| GOBP MICROGLIAL CELL ACTIVATION | 0.00035058 | -1.9052278 | 40 |
| GOBP REGULATION OF MAPK CASCADE | 0.00041326 | -1.464451 | 599 |

| | | | |
|---|---|---|---|
| GOBP MONONUCLEAR CELL DIFFERENTIATION | 0.00041467 | -1.5815887 | 360 |
| GOBP SUPEROXIDE METABOLIC PROCESS | 0.00042285 | -1.8752296 | 56 |
| GOBP ORGANIC ANION TRANSPORT | 0.00043526 | -1.6307288 | 272 |
| GOBP REGULATION OF B CELL MEDIATED IMMUNITY | 0.00045901 | -1.8790796 | 50 |
| GOBP POSITIVE REGULATION OF MAPK CASCADE | 0.00051328 | -1.5444081 | 441 |
| GOBP T CELL ACTIVATION | 0.00052122 | -1.5227882 | 419 |
| GOBP REGULATION OF PHAGOCYTOSIS | 0.00054017 | -1.831916 | 81 |
| GOBP ICOSANOID METABOLIC PROCESS | 0.00059615 | -1.8046337 | 94 |
| GOBP FATTY ACID METABOLIC PROCESS | 0.00060187 | -1.609613 | 326 |
| GOBP POSITIVE REGULATION OF CELL CELL ADHESION | 0.00060769 | -1.6445446 | 255 |
| GOBP CELL KILLING | 0.00061144 | -1.7861556 | 122 |
| GOBP CHEMICAL HOMEOSTASIS | 0.00062846 | -1.3589532 | 959 |
| GOBP HORMONE METABOLIC PROCESS | 0.00062846 | -1.7083877 | 167 |
| GOBP RESPONSE TO METAL ION | 0.00062846 | -1.6214282 | 295 |
| GOBP B CELL RECEPTOR SIGNALING PATHWAY | 0.00076052 | -1.8683047 | 57 |
| GOBP REGULATION OF ANION TRANSPORT | 0.0007892 | -1.4148112 | 714 |
| GOBP REGULATION OF CELL CELL ADHESION | 0.00081787 | -1.5441581 | 389 |
| GOBP ORGANIC ACID TRANSPORT | 0.00082844 | -1.6522277 | 245 |
| GOBP CELL MORPHOGENESIS | 0.00086952 | -1.3635759 | 905 |
| GOBP REGULATION OF MACROPHAGE FUSION | 0.00087509 | -1.6435363 | 5 |
| GOBP REGULATION OF LEUKOCYTE MEDIATED IMMUNITY | 0.00093588 | -1.6990655 | 181 |
| GOBP HETEROTYPIC CELL CELL ADHESION | 0.00099557 | -1.8370475 | 55 |

Table S10. GSEA of HALLMARK pathways of endothelial cells in infected chips with macrophages compared to infected chips without macrophages (adj $P < 0.05$).

| **Gene set** | **Adjusted *P*-value** | **NES** | **Size** |
|---|---|---|---|
| HALLMARK E2F TARGETS | 0.000045 | -1.77957 | 200 |
| HALLMARK G2M CHECKPOINT | 0.000142 | -1.72920 | 199 |
| HALLMARK INTERFERON ALPHA RESPONSE | 0.000165 | 1.821877 | 97 |
| HALLMARK EPITHELIAL MESENCHYMAL TRANSITION | 0.012987 | -1.47110 | 198 |
| HALLMARK MYC TARGETS V1 | 0.012987 | -1.46120 | 199 |
| HALLMARK INTERFERON GAMMA RESPONSE | 0.027272 | 1.451903 | 197 |
| HALLMARK ESTROGEN RESPONSE LATE | 0.032643 | 1.452735 | 190 |
| HALLMARK ALLOGRAFT REJECTION | 0.038769 | 1.398143 | 180 |
| HALLMARK MITOTIC SPINDLE | 0.044924 | -1.32691 | 199 |

Table S11. GOBP analysis of endothelial cells in infected chips with macrophages compared to infected chips without macrophages (adj $P < 0.05$).

| **Gene set** | **Adjusted *P*-value** | **NES** | **Size** |
|---|---|---|---|
| GOBP INNATE IMMUNE RESPONSE | 0.000240 | 1.505022 | 719 |
| GOBP DEFENSE RESPONSE TO OTHER ORGANISM | 0.005446 | 1.420160 | 870 |
| GOBP CHEMICAL HOMEOSTASIS | 0.006837 | 1.384689 | 935 |
| GOBP PATTERN RECOGNITION RECEPTOR SIGNALING PATHWAY | 0.006837 | 1.674636 | 186 |
| GOBP REGULATION OF GLIAL CELL PROLIFERATION | 0.021659 | -2.05548 | 25 |
| GOBP REGULATION OF IMMUNE RESPONSE | 0.021659 | 1.395461 | 732 |
| GOBP POSITIVE REGULATION OF ERK1 AND ERK2 CASCADE | 0.023687 | 1.677957 | 153 |
| GOBP REGULATION OF LYSOSOMAL PROTEIN CATABOLIC PROCESS | 0.023687 | 1.710118 | 8 |
| GOBP TOLL LIKE RECEPTOR SIGNALING PATHWAY | 0.023687 | 1.677879 | 142 |

Table S12. GSEA of KEGG pathways of endothelial cells in infected chips with macrophages compared to infected chips without macrophages (adj $P < 0.05$).

| **Gene set** | **Adjusted *P*-value** | **NES** | **Size** |
|---|---|---|---|
| KEGG COMPLEMENT AND COAGULATION CASCADES | 0.005859 | 1.776323 | 54 |
| KEGG SYSTEMIC LUPUS ERYTHEMATOSUS | 0.005859 | 1.738849 | 84 |

Table S13. Antibody information.

| Antigen | Vendor | Conjugation | Catalog No. | Application |
|---|---|---|---|---|
| CD45 | Thermo Fisher | None | MA1-19111 | Immunofluorescence |
| E-cadherin | BD Biosciences | Alexa555 | 560064 | Immunofluorescence |
| VE-cadherin | Thermo Fisher | Alexa488 | 53-1449-42 | Immunofluorescence |
| Influenza A viral NP | Thermo Fisher | None | MA5-42364 | Immunofluorescence |
| NLRP3 | Thermo Fisher | None | PA5-80857 | Immunofluorescence |
| Caspase-1 | Antibodies Incorporated | FLICA660 | 9122 | Immunofluorescence |
| Anti-mouse IgG | Sigma Aldrich | CF488A | SAB4600035 | Immunofluorescence |
| Anti-rabbit IgG | Sigma Aldrich | CF633 | SAB4600132 | Immunofluorescence |
| DAPI | Sigma Aldrich | N/A | D9542 | Immunofluorescence |
| CD326 | BioLegend | BV421 | 324220 | Flow cytometry |
| CD45 | BioLegend | APC-Fire 750 | 304062 | Flow cytometry |
| Annexin V | BioLegend | FITC | 640906 | Flow cytometry |
| Propidium iodide | Thermo Fisher | N/A | P1304MP | Flow cytometry |
| CD14 | BD Biosciences | BUV395 | 563561 | Flow cytometry |
| CD209 | BD Biosciences | PE-Cy7 | 330114 | Flow cytometry |
| CD163 | BD Biosciences | PE | 556018 | Flow cytometry |

Table S14. Primer information.

| **Name** | **Sequence (5’ to 3’)** | |
|---|---|---|
| GAPDH | Forward | ACATCGCTCAGACACCATG |
| | Reverse | TGTAGTTGAGGTCAATGAAGGG |
| H3N2 viral NP | Forward | TCAAGTGAGAGAAAGTCGGA |
| | Reverse | TCGAAGTCGTAGCCACTGGC |
| CASP1 | Forward | TGAAGGACAAACCGAAGGTG |
| | Reverse | CACATCACAGGAACAGGCATA |
| IL1B | Forward | CAGCCAATCTTCATTGCTCAAG |
| | Reverse | GAACAAGTCATCCTCATTGCC |
| TNF | Forward | TGCACTTTGGAGTGATCGG |
| | Reverse | TCAGCTTGAGGGTTTGCTAC |
| CXCL10 | Forward | GACATATTCTGAGCCTACAGCA |
| | Reverse | CAGTTCTAGAGAGAGGTACTCCT |
| IL6 | Forward | GCAGATGAGTACAAAGTCCTGA |
| | Reverse | TTCTGTTGCCTGCAGCTTC |
| IFNL1 | Forward | GGTTCAAATCTCTGTCACCACA |
| | Reverse | GAAGACAGGAGAGCTGCAAC |
